# Causally informed activity flow models provide mechanistic insight into network-generated cognitive activations

**DOI:** 10.1101/2021.04.16.440226

**Authors:** Ruben Sanchez-Romero, Takuya Ito, Ravi D. Mill, Stephen José Hanson, Michael W. Cole

**Author notes:** Corresponding author (R. Sanchez-Romero). **Data/code availability statement:** Data and code to reproduce our analyses will be publicly available at the project repository upon acceptance of the manuscript. (This information is included in Materials and Methods.). **Ethics statement: We used open access data from the Human Connectome Project (HCP)**, for which all subjects gave signed informed consent in accordance with the protocol approved by the Washington University institutional review board. We abide by the HCP open access use terms and the Rutgers University institutional review board approved use of these data. (This information is included in Materials and Methods.). **Declaration of interests:** The authors have no conflict of interest to declare. **Author statement: Ruben Sanchez-Romero**: Conceptualization, Methodology, Software, Formal Analysis, Writing-original draft, Writing-review & editing, Visualization. **Takuya Ito**: Conceptualization, Methodology, Investigation, Data Curation, Writing-review & editing. **Ravi D. Mill**: Conceptualization, Methodology, Investigation, Data Curation, Writing-review & editing. **Stephen José Hanson**: Conceptualization, Methodology, Resources, Writing-review & editing, Funding acquisition. **Michael W. Cole**: Conceptualization, Methodology, Software, Investigation, Resources, Writing-original draft, Writing-review & editing, Supervision, Funding acquisition.

## Abstract

Brain activity flow models estimate the movement of task-evoked activity over brain connections to help explain network-generated task functionality. Activity flow models have been shown to accurately generate task-evoked brain activations across a wide variety of brain regions and task conditions. However, these models have had limited explanatory power, given known issues with causal interpretations of the standard functional connectivity measures used to parameterize activity flow models. We show here that functional/effective connectivity (FC) measures grounded in causal principles facilitate mechanistic interpretation of activity flow models. We progress from simple to complex FC measures, with each adding algorithmic details reflecting causal principles. This reflects many neuroscientists’ preference for reduced FC measure complexity (to minimize assumptions, minimize compute time, and fully comprehend and easily communicate methodological details), which potentially trades off with causal validity. We start with Pearson correlation (the current field standard) to remain maximally relevant to the field, estimating causal validity across a range of FC measures using simulations and empirical fMRI data. Finally, we apply causal-FC-based activity flow modeling to a dorsolateral prefrontal cortex region (DLPFC), demonstrating distributed causal network mechanisms contributing to its strong activation during a working memory task. Notably, this fully distributed model is able to account for DLPFC working memory effects traditionally thought to rely primarily on within-region (i.e., not distributed) recurrent processes. Together, these results reveal the promise of parameterizing activity flow models using causal FC methods to identify network mechanisms underlying cognitive computations in the human brain.

**Highlights:**

- Activity flow models provide insight into how neurocognitive effects are generated from brain network interactions.
- Functional connectivity methods grounded in statistical causal principles facilitate mechanistic interpretations of task activity flow models.
- Mechanistic activity flow models accurately predict task-evoked neural effects across a wide variety of brain regions and cognitive tasks.

## Introduction

Most traditional explanations of cognitive phenomena are based on focal measures of neural responses to experimental interventions (Cabeza & Nyberg, 2000; Saxe et al., 2006). While this approach has been successful in establishing robust associations between brain regions and cognitive tasks (for example, dorsolateral prefrontal cortex with working memory tasks or fusiform face area with face visual stimuli), these associations by themselves do not explain how task activity in a brain region is generated from underlying causal processes (e.g., brain network interactions). Patterns of task-related neural activations have also been used in multivariate analyses to explain differences between task conditions (Kriegeskorte et al., 2008; Norman et al., 2006), but also fail to mechanistically explain how activations are generated from underlying causal processes. Another strategy to characterize associations between cognitive tasks and brain regions is to analyze changes in inter- or intra-region connectivity due to task manipulations (Gordon et al., 2014; Jolles et al., 2013; Vatansever et al., 2017). However, these studies do not typically assess the role of task-related neural activations, which are more clearly linked to cognition and behavior (e.g., activations in M1 are known to cause motor responses). Other explanations of cognitive effects are based on artificial neural network models that try to reproduce empirically observed cognitive responses (Thomas & McClelland, 2008). However, most of these models are not constrained by empirical brain data (biological network architectures or signals), and thus provide only limited mechanistic insight. While each approach contributes important information regarding the role of brain processes in behavior and cognition, these approaches do not provide empirically-supported neural explanations of how behavioral and cognitive phenomena are causally generated.

Developed by considering the limitations of these strategies to explain cognitive phenomena, the activity flow mapping (actflow) framework (Cole et al., 2016) uses empirical brain data to produce a generative connectionist model of task-related activations. The resulting model integrates empirically-derived functional/effective connectivity (FC) networks and empirical task activations (**Figures 1F**-**G**) to provide data-driven mechanistic insight into network-supported cognitive processes in the human brain.

**Figure 1.**
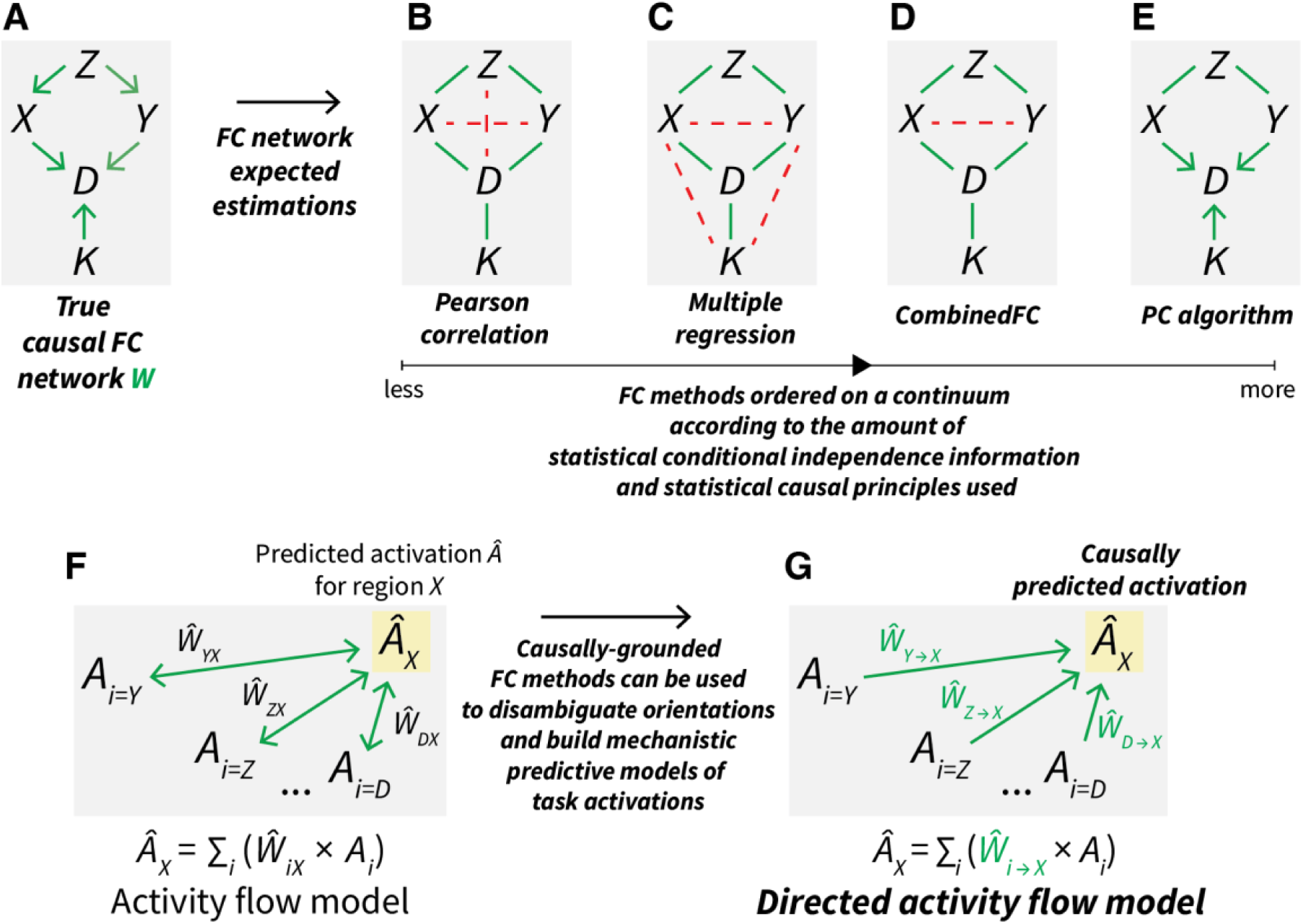
Causally-grounded functional connectivity methods can be used to build directed activity flow models. (A) An example of a true causal FC network, denoted as W, for five neural regions time series *Z*, *X*, *Y*, *D* and *K*. The green arrows represent direct causal functional associations between the time series. **(B)** The expected network when using Pearson correlation FC to recover the true mechanism from panel B. Green lines indicate correctly inferred undirected connections, while red dashed lines indicate incorrectly inferred connections. Incorrectly inferred connections resulted from not controlling for one causal confounder (*X* ← *Z* → *Y*) and two causal chains (*Z* → *X* → *D* and *Z* → *Y* → *D*). **(C)** The expected network when using multiple regression FC. The association between each pair of time series is conditioned on the rest of the regions to control for confounders and chains. In this case, the incorrectly inferred connections resulted from three conditioned-on causal colliders (*X* → *D* ← *Y*, *X* → *D* ← *K* and *Y* → *D* ← *K*). **(D)** The network recovered by combinedFC. Thanks to its zero-correlation check—based on the statistical behavior of colliders (see CombinedFC section)—combinedFC removed two of the spurious connections from conditioned-on colliders. Nevertheless, for this mechanism the zero-correlation check cannot remove the remaining spurious connection because the confounder (*X* ← *Z* → *Y*) forces a non-zero correlation between *X* and *Y*. **(E)** The directed network recovered by the PC algorithm. By iteratively testing associations with conditioning sets of increasing size and then applying a series of orientation rules, PC can infer the true FC mechanism from panel C, with the exception of the direction of two connections (see the PC algorithm section for details on why some connections cannot be oriented by this method). **(F)** Activity flow predictive model. The inferred task-evoked activation *Â* of a held-out neural region *X* (yellow box) is predicted as a linear function of the inferred FC weights (*Ŵ*, green bidirectional arrows) and the actual task-evoked activations *A* from the rest of the *i* connected regions. The bidirectional arrows reflect the ambiguity of the inferred FC with respect to the true causal orientation. **(G)** Directed activity flow model using data-driven information about the causal direction of the functional connections. The task-evoked activation *Â* of a neural region *X* (yellow box) is predicted as a linear function of the causal FC weights (*Ŵ*, green unidirectional arrows) and the actual task-evoked activations *A* from the *i* causal source regions. The unidirectional arrows convey that directed FC methods can be helpful to infer the causal direction of connections.

The actflow framework has been previously applied in a variety of studies to investigate: the flow of task-related activity via whole-brain resting-state networks (Cole et al., 2016); the fine-scale transfer of information-representing task activity between specific pairs of functionally connected regions (Ito et al., 2017); the relevance of task-state functional networks in communicating task-related neural responses (Cole et al., 2021); the disruption of task activations from altered functional networks in pre-clinical Alzheimer’s disease (Mill et al., 2020); the disruption of task activations from altered activity flows in schizophrenia (Hearne et al., 2021); the role of specific brain networks in a visual shape completion task (Keane et al., 2021); and the cortical heterogeneity of localized and distributed cognitive processes (Ito, Hearne, & Cole, 2020). These studies evidence that actflow modeling can successfully provide insights about the underlying network processes generating the observed neural responses caused by exogenous cognitive task manipulations.

In the aforementioned studies, actflow models parameterized with FC networks estimated with Pearson correlation (pairwise association; the field standard) or with multiple regression (fully conditional associations) provided accurate predictions of task-evoked activations, suggesting that these methods can capture to some degree, relevant properties of the mechanisms supporting task-related functionality. Nevertheless, these FC methods pose limitations for the interpretability and intervention potential of actflow models. For example, they are inherently undirected, thus from their results we cannot make inferences about the causal direction of the activity flow evoked by a task manipulation. In addition, these FC methods are prone to false-positive inferences in the presence of uncontrolled common causes (causal confounders, *X* ← *Z* → *Y*), uncontrolled causal chains (*X* → *Z* → *Y*), and incorrectly controlled common effects (causal colliders, *X* → *Z* ← *Y*) (**Table 1**) (Reid et al., 2019). We hypothesize that these false-positive connections can bias actflow predictions by producing incorrect estimations of the true underlying functional networks supporting task computations.

**Table 1.** Ability of FC methods to account for the presence of basic causal structures and to infer orientations. When these causal structures are not accounted for, FC methods will infer false-positive connections. For example, in the presence of a causal confounder, Pearson correlation will infer a false-positive connection *X* — *Y* since it does not account for the confounding effect of region *Z*.

| | Causal<br>confounder<br>$X \leftarrow Z \rightarrow Y$ | Causal<br>chain<br>$X \rightarrow Z \rightarrow Y$ | Causal<br>collider<br>$X \rightarrow Z \leftarrow Y$ | Causal confounder &<br>collider<br>$X \leftarrow Z \rightarrow Y \text{ \& } X \rightarrow D \leftarrow Y$ | Orientations<br>inferred |
| --- | --- | --- | --- | --- | --- |
| **Pearson correlation** | Does not account for | Does not account for | Accounts for | Does not account for | No |
| **Multiple regression** | Accounts for | Accounts for | Does not account for | Does not account for | No |
| **CombinedFC** | Accounts for | Accounts for | Accounts for | Does not account for | No |
| **PC algorithm** | Accounts for | Accounts for | Accounts for | Accounts for | Some |

To overcome these limitations, we propose building actflow models parameterized with FC methods that more effectively use statistical conditional independence information and statistical causal principles to reduce the risk of false-positive connections and infer (when possible) the direction of connections. In an ideal biologically-constrained mechanistic actflow model, task-related activations of a target neural region would be predicted from the actual activations of only its true direct causal source regions (**Figure 1H**). Pursuing this kind of mechanistic model implies that we are not only interested in predicting neural activations but also in hypotheses about how a target neural region may react to exogenous (e.g., experimental interventions using transcranial stimulation) or endogenous (e.g., age-related cognitive decline) changes in its afferent region activations, connectivity patterns or both. This interest in mechanism reflects both a desire for fundamental causal understanding of the brain as a complex system, as well as the practical desire to develop causal interventions to cure (or reduce the burden of) brain diseases (i.e., treatments).

We first test our hypothesis by evaluating the impact of different FC methods on actflow model prediction accuracy in simulations and empirical data. We order FC methods on a continuum according to the complexity of each method’s underlying algorithm, defined by the amount of statistical conditional independence information and statistical causal principles used. This reflects the desire for many neuroscientists to use the simplest method possible—due to complexity adding more features needing validation/justification while also making it more difficult to communicate findings. This can be thought of as a kind of Occam’s Razor principle, in which the cost of additional methodological complexity must be justified by substantially better results in terms of empirical validity and/or theoretical insight. We start with the field-standard Pearson pairwise correlation (no use of conditional independence information), followed by multiple regression (use of fully conditional independence information), then combinedFC (Sanchez-Romero & Cole, 2021) (use of fully conditional independence information and causal principles to detect the presence of causal colliders), and finishing with the Bayesian network-based PC (Peter-Clark) algorithm (Spirtes et al., 2000) (iterative use of increasing size sets of conditional independence information and use of causal principles to detect causal colliders and orient connections) (**Figures 1A**-**E** and **Table 1**). We use the PC algorithm as we consider it a tractable example of how causal principles can be used to effectively integrate pairwise and conditional independence information and also orient connections (see PC algorithm section, **Figure 1E**, **Table 1** and limitations in Discussion).

While we expect that FC methods leveraging more conditional independence information and causal principles will produce better actflow predictions (by reducing the deleterious effect of spurious pathways), we are aware that not controlling for confounders can still produce models with prediction potential. In principle, we could predict information from region *Y* using information from region *X* even if *X* is not a cause of *Y* (or vice versa), as long as they are confounded by a third region. This suggests the possibility of FC models with good prediction but spurious causal information (a spurious connection between *X* and *Y* in this example). Critically, however, even if prediction accuracy were lower (but above chance) for an FC method grounded in statistical causal principles, this study’s results would support its practical utility (i.e., for predicting empirical effects) in addition to its theoretical utility (i.e., improving causal interpretability of the resulting model). This reflects our joint goals of using actflow to produce predictive models that generate neurocognitive processes of interest (e.g., fusiform face area activity during face perception) while also being grounded in statistical causal principles.

Finally, we illustrate with empirical data how actflow model-based explanations can go beyond isolated accounts of localized neural activation, or isolated accounts of neural connectivity patterns, by providing mechanistic insight into the network processes that generate the observed neural responses caused by cognitive task manipulations. Specifically, we build a directed actflow model focusing on a single region of the DLPFC. This model is parameterized with PC-algorithm FC, with the goal of identifying likely network interactions contributing to n-back working memory task activity. Notably, traditional accounts of DLPFC activity during working memory tasks focuses on persistent activity in DLPFC (Curtis & D’Esposito, 2003; Sreenivasan et al., 2014), which is typically thought to result from within-region recurrence (Wang et al., 2013). In contrast, actflow models rely exclusively on activity from distal brain regions, such that an actflow model accurately generating DLPFC activity during a working memory task would provide evidence for an alternate (or complementary) distributed account of working-memory-related DLPFC activity.

Confirmation that FC methods that leverage statistical conditional independence information and statistical causal principles lead to accurate actflow predictions would demonstrate the utility of using actflow models from brain data to develop mechanistic explanations of cognitive functions from exogenous task manipulations. The mechanistic insight provided by actflow models represents a starting point for what could be considered a full neurocognitive mechanistic explanation, which would identify the full chain of effects from exogenous stimulus presentation to network-supported cognitive neural activations to behavioral response (Ito, Hearne, Mill, et al., 2020; Weichwald & Peters, 2021).

## Materials and Methods

### Activity flow mapping

Activity flow mapping (actflow) is a predictive model to explain local task-related neural activations as the product of task-evoked activity flowing through pathways of functional brain connections (Cole et al., 2016). Formally, for a set of measured brain regions **V**, the task-related activation *A_X_* for brain region *X*, can be expressed as *A_X_*= *f*(W*_X_*, A**_V_**_\{*X*}_), where W*_X_* are the connections of *X* with the rest of the regions, A**_V_**_\{*X*}_ are the activations of all regions in **V** except *X*, and *f*() is a function relating connections and activations. Following (Cole et al., 2016), we assume *f*() is a linear function and implement the actflow model for a particular held-out region as *Â_X_* = Σ*_i_*_∈**V**\{*X*}_*Ŵ_iX_A_i_*, where the predicted activation (*Â_X_*) is the sum of the actual activations of all other regions (*A_i_*), weighted by their estimated connectivity values with *X* (*Ŵ_iX_*) (**Figure 1F**). (This function corresponds to a standard neural network linear propagation and activation rule (Hanson & Burr, 1990; Rumelhart et al., 1986).)

Theoretically, the Markov blanket of a target variable is the set of its direct causes, direct effects and other direct causes of those direct effects. This set of variables accounts for all relevant information to optimally predict the target variable (Guyon et al., 2008). The above definition of actflow does not differentiate between direct causes and direct effects in the connectivity pattern, and predicts using all inferred connected regions. By using all connected regions to predict the activation of a particular held-out region we are leveraging information from its direct causes (as the linear combination of the afferent (incoming) connection weights and the source activations), and from its direct effects (as the linear combination of the efferent (outgoing) connection weights and the effect activations). This suggests that an actflow model can be parameterized with an undirected FC method (that does not differentiate causes and effects) and achieve good prediction accuracy, since it is effectively using a subset of the Markov blanket (specifically, direct causes and direct effects). In our analysis we compare actflow models parameterized with undirected FC networks derived from: correlation, multiple regression, combinedFC and the undirected output of PC (see PC section).

Despite the likelihood that using all truly connected regions is optimal for predicting task-evoked activations, we want to impose a causal biological constraint on the actflow models, such that the activation of a held-out target region is predicted from the actual task-evoked activations of its direct causal sources and the corresponding afferent connection weights (**Figure 1G**). From a mechanistic perspective, our goal is to hypothesize how the activation of a target region may react to changes in its direct-cause activations. Nevertheless, moving towards this kind of biologically-constrained predictive model comes with a potential reduction in predictive power, since we will predict a target region only using its direct causes, not leveraging the full Markov blanket.

With this idea in mind, we define a biologically-constrained mechanistic linear actflow model as *Â_X_* = Σ*_i_*_∈**V**\{*X*}_*Ŵ_i → X_A_i_*, where for a set of measured regions **V**, *Ŵ_i → X_*are the estimated directed connections from direct sources *i* to held-out target region *X*. In contrast to the first actflow definition (**Figure 1F**), this model uses only the estimated direct sources to predict the task-evoked activations for a target region (**Figure 1G**). The challenge with this actflow mechanistic model is that to obtain connectivity estimates *Ŵ_i → X_*, we necessarily need a directed FC method. Here, we use the PC algorithm to estimate the required directed networks.

Finally, as in Cole et al. (2016), we measured the prediction accuracy of actflow models using the Pearson correlation *r* between predicted and actual activations, and compared it across the different FC methods used to parameterize the models. Activity flow mapping prediction and evaluation analyses were performed with the Python open-source Actflow Toolbox (available at https://colelab.github.io/ActflowToolbox).

### Functional connectivity methods

Here, we ordered functional connectivity methods in a continuum depending on the amount of statistical conditional independence information and statistical causal principles used, and focus on its limitations regarding risk of inferring false-positive connections and capability to infer direction of connections.

#### Correlation

We start with pairwise associative methods, such as Pearson correlation or mutual information (a way to measure non-linear statistical associations). These methods do not use conditional independence information and do not hold causal assumptions about the generating mechanism giving rise to the observed association. For example, a non-zero Pearson correlation between the time series of two brain regions *X* and *Y* indicates a functional association between these regions, but no further knowledge about the nature of this association can be derived from it. We cannot conclude if the observed non-zero pairwise correlation resulted from a causal mechanism where one brain region is the direct cause of the other (*X* → *Y* or *X* ← *Y*), or one region is the indirect cause of the other (causal chain, *X* → *Z* → *Y*), or a third region is a common cause of the two regions (causal confounder, *X* ← *Z* → *Y*) (Reichenbach, 1956), or a combination of these cases. This ambiguity impedes mechanistic interpretations of correlation-based network connections. In practice, pairwise associative methods do not control for the effects of causal confounders and chains, thus inferring false-positive connections in the estimated FC network (**Figure 1B** and **Table 1**). In linear activity flow models, these false-positive connections will create false pathways through which task activity will be incorrectly added, biasing the prediction of the task-evoked activations.

We applied Pearson correlation *r_XY_* = *cov*(*X*,*Y*)/(*std*(*X*)*std*(*Y*)), where *X* and *Y* are time series for two brain regions, *cov*() is the sample covariance and *std*() is the sample standard deviation; and used a two-sided *z*-test with significance threshold of *p*-value < α = 0.01, for every individual simulation and empirical dataset.

#### Multiple regression

Multiple regression is used to compute the statistical association between one region time series and every member of a set of regressor time series—in FC analysis this set is usually the rest of the brain regions in the dataset—where each association is conditioned on the rest of the regressors (fully conditional association). Conditioning on the rest of the regions in the dataset controls for false-positive connections arising from the effect of causal chains and causal confounders. Despite controlling for this type of false-positive connections, a fundamental limitation of multiple regression as an FC method is that by fully conditioning on the rest of the brain regions it may infer a false-positive association between two unconnected regions if these two regions are causes of a third one (collider, *X* → *Z* ← *Y*) (**Figure 1C** and **Table 1**) (Berkson, 1946; Bishop, 2006; Kiiveri et al., 1984; Reid et al., 2019). The presence of colliders in a causal structure—which we cannot tell in advance—implies that any connection inferred by multiple regression could in principle be a false-positive connection. For example, Sanchez-Romero & Cole (2021) showed that in simulated networks with a larger proportion of colliders relative to confounders, multiple regression returns a higher number of false-positive connections than correlation.

We applied ordinary least squares linear multiple regression *Y* = ***β***_0_+***β***_1_*X*_1_+***β***_2_*X*_2_+…+***β****_k_X_k_*+*e_Y_*, where *Y* is the time series for a brain region, *X*_1_ to *X_k_* are the time series for the rest of the regions in the dataset, ***β***_1_ to ***β****_k_* are the corresponding regression coefficients, ***β***_0_ is the intercept and *e_Y_*is the regression error of *Y*. We used a two-sided *t*-test for the regression coefficients with significance threshold of *p*-value < α = 0.01, for every individual simulation and empirical dataset.

#### CombinedFC

The combinedFC method (Sanchez-Romero & Cole, 2021) proposes a causally-principled solution to avoid false-positive connections from conditioning on colliders. The strategy of combinedFC is based on the observation that for a collider *X* → *Z* ← *Y*, the pairwise correlation of the two causes *X* and *Y* will be zero; while the multiple regression *X* = *aY* + *bZ* (or *Y* = *aX* + *bZ*), where the common effect *Z* is being conditioned on, will infer a non-zero regression coefficient a between *X* and *Y*.

CombinedFC leverages this observation to detect and remove false-positive connections from conditioning on colliders. In a first step, the method computes the multiple regression for each brain region on the rest of the regions in the dataset. In a second step, it checks for each non-zero multiple regression coefficient if its corresponding pairwise correlation is zero. If this is the case, there is evidence of a false-positive connection from conditioning on a collider and combinedFC removes this connection from the network (**Figure 1D** and **Table 1**). Using combinedFC we have more evidence to conclude that the inferred network does not include false-positive connections from unconditioned causal confounders or chains—thanks to the initial multiple regression fully conditioning—or false-positive connections from conditioning on colliders—thanks to the zero-correlation check. Nonetheless, Sanchez-Romero & Cole (2021) have shown that in the presence of certain challenging causal patterns, for example a mix of a causal confounder and a collider (e.g., **Figure 1A** and **Table 1**), combinedFC will inevitably produce false-positive connections (**Figure 1D** and **Table 1**). It is the risk of these false-positives that limits combinedFC to unequivocally orient detected colliders. Thus, as correlation and multiple regression, combinedFC returns an undirected network, albeit with less risk of false-positive connections.

Here, we implement combinedFC with two modifications. In the first step of combinedFC, instead of using linear multiple regression to evaluate conditional associations, we used partial correlation using the inverse of the covariance matrix, which is a faster and equivalent way to determine significant connections. The second modification is considering the output of combinedFC as an initial feature selection step (Guyon et al., 2008; Guyon & Elisseeff, 2003), and compute the final FC weights by regressing each region only on its connected regions (selected features) in the combinedFC network. We have seen that this second modification produces FC weights that result in more accurate activity flow predictions than the original weights of combinedFC.

For every individual simulation and empirical dataset, we applied combinedFC with a two-sided *z*-test and significance threshold of *p*-value < α = 0.01 for the partial correlation, and of *p*-value > α = 0.01 for the zero-correlation check (following Sanchez-Romero & Cole (2021)). After the feature selection, we computed FC weights with linear multiple regression and no significance test.

#### PC algorithm

The Peter-Clark (PC) algorithm (Spirtes et al., 2000; Spirtes & Glymour, 1991) provides a discovery strategy that overcomes the limitations of the three methods described above. It controls for false-positive connections created by casual chains, confounders and conditioned-on colliders, even in challenging causal patterns (**Figure 1E** and **Table 1**). In addition, after undirected connections are estimated (adjacency discovery phase), the PC algorithm applies a series of orientation rules to infer, when possible, the causal direction of connections (orientation discovery phase). The outcome of the algorithm is a network of undirected and directed connections from which a set of equivalent (i.e., encode the same conditional independence information from the data) fully directed networks can be read out. The set of equivalent fully directed networks is built by listing all the possible combinations of orienting undirected connections (if any), as long as no new colliders are created. For example, in **Figure 1E** the set of three equivalent fully directed networks is built by orienting the undirected connections *X* — *Z* — *Y*, as *X* ← *Z* → *Y*, *X* → *Z* → *Y* and *X* ← *Z* ← *Y*. Importantly, without additional information (e.g., from experimental causal interventions) we are not able to disambiguate which of the three is the true directed network (Eberhardt et al., 2005). (In the Bayes networks literature, this equivalence set is referred to as the Markov equivalence class (Verma & Pearl, 1992).)

We used an order-independent version of the PC adjacency discovery phase known as PC-stable (Colombo & Maathuis, 2014; termed FAS-stable in Sanchez-Romero et al., 2019), and include pseudocode in **Box 1 (1)**. In short, PC starts with a fully connected undirected network and checks for every pair of brain regions if they are correlated or not. If two regions are not correlated, PC removes their adjacency from the network. Next, the algorithm checks for every pair of still connected regions if they are correlated conditioning on one other region. (This implies a conditional correlation with a conditioning set of size one. In the pseudocode, the size of the conditioning set **S** is referred to as *depth* (**Box 1 (1).3**).) If two regions are not conditionally correlated on one other region, PC removes their connection from the network. For every pair of still connected regions, PC keeps testing correlations conditioning on two other regions (conditioning set of size two), three other regions and so forth, until no more connections can be removed from the network. Note that when PC evaluates conditional correlations, it does it iteratively through all possible combinations of conditioning sets of size one, two, three and so forth, until it finds, if any, a set **S** that makes the regions not conditionally correlated (**Box 1 (1).3.b**).

The implementation we used of PC computes the conditional correlation for any two regions *X* and *Y* conditioning on a set **S** (**Box 1 (1).3.b.i**), using the inverse of the covariance matrix (precision matrix P) for *X*, *Y* and **S**, to obtain the conditional correlation coefficient *r_XY_*_|**S**_ = -P*_XY_*/*sqrt*(P*_XX_*P*_YY_*), where *sqrt*() is the square root function and P*_XY_* is the entry for *X* and *Y* in the precision matrix. To determine statistical significance, first the conditional correlation coefficient *r_XY_*_|**S**_ is transformed to a Fisher *z* statistic *ƶ* = *tanh*^-1^(*r_XY_*_|**S**_)*sqrt*(*N*−|**S**|−3), where *N* is the number of datapoints and |**S**| is the size of the conditioning set (number of regions conditioned on); then, for a two-sided *z*-test, we compute the *p*-value = 2(1−*cdf*(*abs*(*ƶ*)), where *abs*() is the absolute value and *cdf*() is the cumulative distribution function for a standard normal distribution. For a user-chosen α significance threshold, if *p*-value > α, then we conclude that regions *X* and *Y* are not correlated conditioning on the set of regions **S**. (In the PC algorithm this result implies removing the network connection between *X* and *Y*.) A significance threshold of α = 0.01 was set for all applications of PC.

It is important to note that we implement the PC algorithm with conditional correlations to estimate the required conditional associations but other approaches can be used, such as conditional mutual information, or other non-linear, non-Gaussian, conditional association measures (Ramsey, 2014; Zhang et al., 2011), depending on the properties of the distributions and functional associations of the data under study.

###### Box 1.

PC algorithm pseudocode

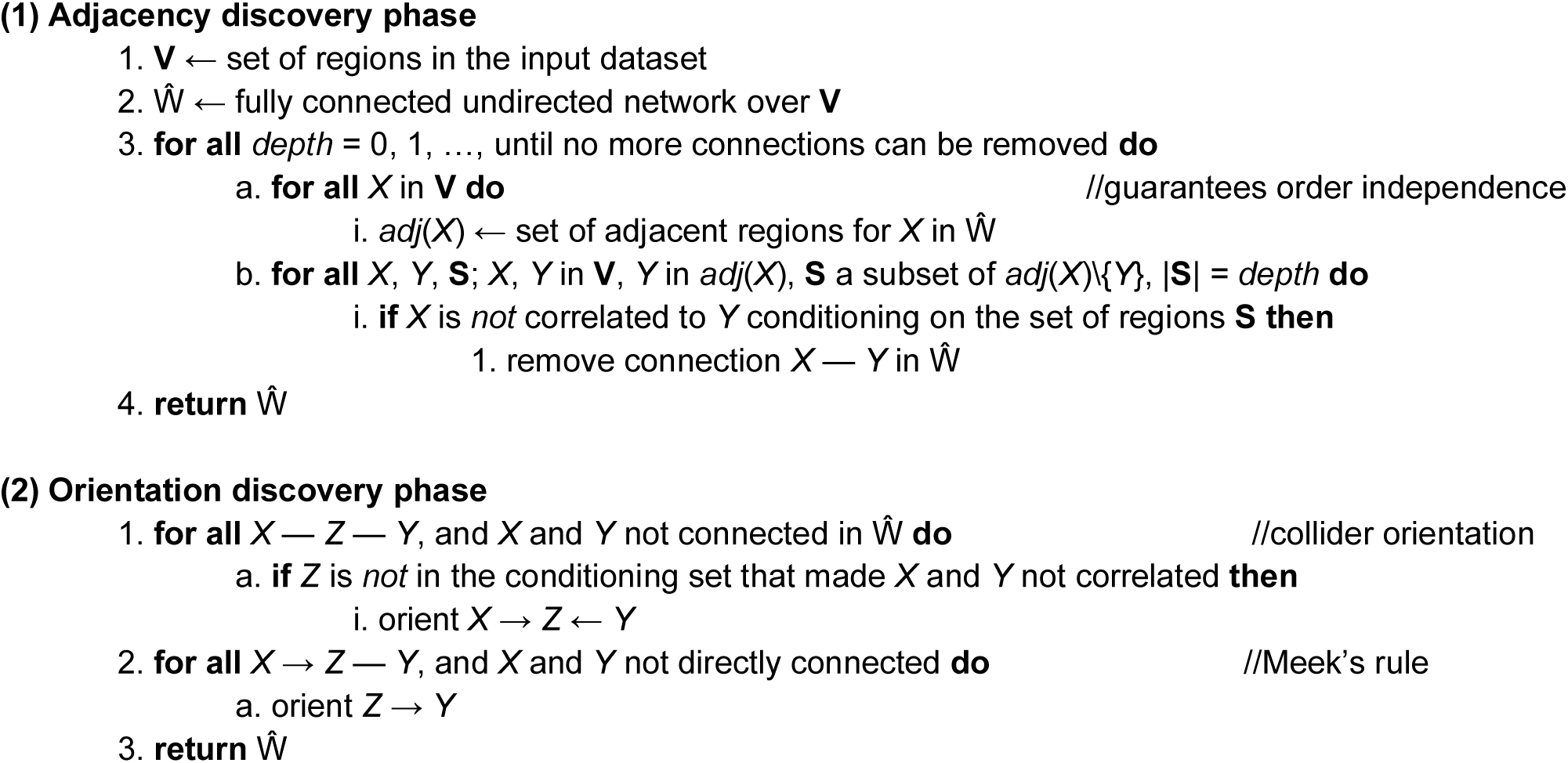

The second phase of our implementation of the PC algorithm applies two rules to orient, when possible, the adjacencies from the first phase. We include the orientation discovery phase rules in **Box 1 (2)**. The first rule is based on causal principles about conditional independencies implied by collider structures and is part of the original implementation of PC (Spirtes & Glymour, 1991). The collider orientation rule states that if in a network, a region *X* is adjacent to a region *Z*, and *Z* is adjacent to a region *Y*, and *X* and *Y* are not adjacent (triple, *X* — *Z* — *Y*), if *Z* is not in the conditioning set that made *X* and *Y* not correlated, then necessarily these regions form a collider *X* → *Z* ← *Y* (**Box 1 (2).1**). If the opposite were true, and *Z* were in the conditioning set that made *X* and *Y* not correlated, then we would not be able to orient this triple, because the three other possibilities *X* → *Z* → *Y*, *X* ← *Z* ← *Y* or *X* ← *Z* → *Y*, equally imply that *X* is not correlated with *Y* conditioning on *Z*. In this particular case, only collider structures produce unambiguous conditional correlations that can be used to orient adjacencies (see **Figure 1E** for an example).

In practice, when the fMRI time series have a small number of datapoints, some conditional correlation estimates may be inaccurate and we could end up incorrectly orienting colliders. To minimize this risk, Ramsey (2016) implemented the collider orientation rule with the max *p*-value heuristic. The heuristic consists in computing the *p*-values of the correlations of regions *X* and *Y* conditioning on every possible subset of regions adjacent to *X*. Then, choosing the conditioning subset corresponding to the maximum *p*-value, and if region *Z* is not in this subset, orienting the triple as a collider *X* → *Z* ← *Y*. By choosing the conditioning subset with the maximum *p*-value we try to guarantee that from all the possible subsets, we select the one that optimally assures that *X* and *Y* are conditionally not correlated.

Importantly, Ramsey (2016) also showed that with inaccurate conditional correlation estimates due to small number of datapoints, we may end up orienting two conflicting colliders in the network. For example, for two triples we could conclude *X* → *Z* ← *Y* and *Z* → *Y* ← *D*, which implies a conflicting orientation for *Z* and *Y*. The problem is deciding which collider orientation we should remove. Ramsey (2016), following the same ideas of the max *p*-value heuristic, suggests sorting all the previously inferred colliders from high to low according to its *p*-value—from the max *p*-value heuristic—and remove a collider orientation if it conflicts with any higher *p*-value collider. Ramsey (2016) showed in simulations the improvement in orientation accuracy from these two heuristics, so in our implementation of PC we used the collider orientation rule with the max *p*-value heuristic followed by the collider conflict resolution heuristic.

Meek (1995) introduced a set of orientation rules that in some cases can complement the collider orientations. This second orientation rule (**Box 1 (2).2**) is based on the assumption that the collider orientation rule properly detected all existing colliders in the network, such that no new colliders are allowed. Thus, for *X* → *Z* — *Y* we can orient *Z* → *Y*, since the opposite direction *Z* ← *Y* will create a new collider, and that is not allowed. The rest of Meek’s rules leverage the assumption that the underlying causal network does not contain cycles, and thus orient adjacencies avoiding the formation of cycles. Since we know the brain contains feedforward and feedback structures supporting communication between regions, the assumption of no cycles is incorrect in this case. For this reason, we did not implement those orientation rules and instead retained undirected connections that may suggest the presence of cycles. For mechanistic interpretations of actflow models, we consider it more problematic to orient a connection in the incorrect causal direction than to provide no orientation at all.

The output of the PC algorithm is an unweighted network Ŵ from which connection weights can be estimated. Using Ŵ as a starting point we derived two different FC approaches. In the first, for each region *X*, we get *Pa*(*X*) the set of causal sources (parents) of *X* in network Ŵ, and solve the linear regression *X* = ***β****_Pa_*_(*X*)_*Pa*(*X*). The elements of the estimated vector of regression coefficients ***β****_Pa_*_(*X*)_ are considered the weights for the source connections into *X*. For example, in *X* → *Z* ← *Y*, *Z* = ***β****_Pa_*_(*Z*)_*Pa*(*Z*) = ***β****_XZ_X* + ***β****_YZ_Y*, such that the estimate for ***β****_XZ_* is the weight for the directed connection *X* → *Z*, and equivalently for ***β****_YZ_*. Doing this for every region outputs a FC network, where each directed connection *X* → *Y* represents a causal hypothesis, in the sense that, keeping all other regions fixed, a change of one unit in *X* will cause an expected change of ***β****_XY_* in *Y* (Pearl, 2000; Spirtes et al., 2000; Woodward, 2005). In our biologically-constraint mechanistic actflow model, using a directed FC network implies predicting task-related activity for a held-out target region using only its putative causal sources (**Figure 1E**). Hereafter we refer to this FC method simply as the PC algorithm or PC.

As mentioned above, the network output by PC encodes a set of equivalent fully directed networks. In practice, as we increase the number of neural regions analyzed, the size of this set grows exponentially (He et al., 2015), making it too expensive (or infeasible) to evaluate all possible equivalent networks across simulations and empirical data. In this first FC approach, to reduce the computational cost we will only consider the original directed connections of the PC output network and not all the possible oriented connections included in the equivalence set. Thus, we simply remove undirected edges from the graphs resulting from the PC algorithm.

Our second FC approach is motivated by 1) the assumption that accurately predicting the activation of a held-out target region can improve by using information from both its true direct causal sources and its true direct causal effects (a subset of its Markov blanket) (Aliferis et al., 2010; Fu & Desmarais, 2010), and 2) an interest in comparing the PC adjacency network to the networks inferred by the other undirected FC methods tested here. In this second approach, for each region *X*, we get *adj*(*X*), the set of adjacent regions for *X* in network Ŵ, and solve the linear regression *X* = ***β****_adj_*_(*X*)_*adj*(*X*). The elements of the estimated vector of regression coefficients ***β****_adj_*_(*X*)_ are considered the connectivity weights for the adjacencies of *X*. For example, in *X* → *Y* → *Z*, *Y* = ***β****_adj_*_(*Y*)_*adj*(*Y*) = ***β****_YX_X* + ***β****_YZ_Z*. Essentially, we are computing the FC weights using every adjacent region, which may include, depending on the inferred causal pattern, only direct causal sources, only direct causal effects or a combination of both—as in the example. These FC weights disregard the orientation of the connections and thus no longer have as straightforward a mechanistic interpretation as in the above PC method. Hereafter we refer to this FC method as PC-adjacencies or PCadj.

Our implementation of the PC algorithm (with the removal of certain Meek orientation rules as described above) is available at [repository available upon acceptance of the manuscript], and it is a Python wrapper of the PC algorithm from the Java open-source Tetrad software version 6.7.1 (available at https://github.com/cmu-phil/tetrad).

### Simulated causal networks and data

As described above, activity flow analysis requires FC estimates from resting-state data and task-evoked activations from task-state data. Our general simulation strategy consisted in first creating a synthetic ground-truth causal network (directed graph), parameterizing and instantiating functional interactions to create a resting-state network and associated dataset, and then simulating a task-state network by introducing small random modifications to the original resting-state network coefficients (to simulate observed task-related deviations from resting-state connectivity (Cole et al., 2021; Ito, Brincat, Siegel, et al., 2020)), plus an exogenous task input variable feeding into the network to produce a task-state dataset.

Simulation of resting-state networks and data closely follows Sanchez-Romero & Cole (2021). Ground-truth resting-state networks were based on a directed random graphical model that has a preference for common causes and causal chains than for colliders, and includes two-node and three-node cycles. All networks were simulated with 200 nodes and an average connectivity density of 5% (percentage of connections out of total possible). The network connectivity coefficients were instantiated by sampling from a uniform distribution in the interval [0.1, 0.4), and randomly setting 10% of the coefficients to its negative value. (This proportion was chosen to approximate the proportion of negative functional connections (15%) observed in the empirical fMRI analysis of Sanchez-Romero & Cole (2021).) To generate resting-state data we used a general causal linear model X = WX+E, where X is a dataset of nodes (nodes × datapoints), W a directed network encoded as a matrix of connectivity coefficients (nodes × nodes), with direction going from column to row, and E a set of independent noise terms (intrinsic activity) (nodes × datapoints). We simulated 1000 datapoints for resting-state data X by solving this model for X = (I-W)^-1^E, where I is the identity matrix (nodes × nodes), W the simulated resting-state network, and pseudo-empirical datapoints for E (Sanchez-Romero & Cole, 2021) were instantiated by randomizing preprocessed fMRI resting-state data across datapoints, regions and participants, from the Human Connectome Project (HCP). Using pseudo-empirical terms E allowed us to simulate resting-state data X that better capture some of the distributional properties of our empirical fMRI.

To create a task-state networks W*_T_* whose connectivity coefficients reflect deviations from rest, we took the previously simulated resting-state network W and defined each task-state coefficient, with equal probability, as: (a) one standard deviation above the corresponding resting-state coefficient, or (b) one standard deviation below, or (c) equal to the resting-state coefficient. (The standard deviation value corresponds to that of the positive connectivity coefficients in resting-state network W.) To generate task-state data we first defined one exogenous task input continuous variable *T* (simulating one task condition) with pseudo-empirical randomized BOLD datapoints from the HCP dataset (defined as above) (1 × datapoints). Then, we defined a task connectivity vector C (nodes × 1), that specifies the network nodes *directly* affected by the task variable *T*. We randomly chose 10% of nodes to be directly affected by *T*, and sampled task connectivity coefficients from a uniform distribution in the interval [0.1, 0.4). (Note that other nodes can also be affected by *T* but through indirect causal paths.) Expanding the causal linear model to include the task-related elements, we now have X*_T_* = W*_T_*X*_T_*+C*T*+E*_T_*. We simulated 1000 datapoints for task-state dataset X*_T_* by solving the model for X*_T_* = (I-W*_T_*) ^-1^(C*T*+E*_T_*). As with the resting-state data we used pseudo-empirical datapoints for E*_T_*.

Finally, simulated task-evoked activations were estimated individually for each node *X_T_* by a standard fMRI general linear model. In other words, a linear regression of the form *X_T_* = *aT*, where the estimated coefficient *a* represents the task-evoked activation, and reflects direct and indirect causal effects of the task variable *T* on the node *X_T_*.

200 instantiations of the simulated models were generated to compare (1) the accuracy of the different FC methods to recover resting-state networks, and (2) the prediction accuracy of the activity flow models parameterized with these networks. Analyses were run in the Rutgers-Newark high-performance computing cluster AmarelN (https://oarc.rutgers.edu/resources/amarel), using one node, 2 cores, and 64G RAM.

### Empirical fMRI data

Resting and task-state empirical fMRI data were used to compare the accuracy of activity flow predictions under the five different FC methods tested here. We used open access fMRI resting and task-state data from a subset of 176 participants from the minimally-preprocessed HCP 1200 release (Glasser et al., 2013; Ugurbil et al., 2013; Van Essen et al., 2013). All subjects gave signed informed consent in accordance with the protocol approved by the Washington University institutional review board. We abide by the HCP open access use terms and the Rutgers University institutional review board approved use of these data. These participants were selected by passing the following exclusion criteria as described in Ito et al. (2020): anatomical anomalies found in T1w or T2w scans; segmentation or surface errors as output from the HCP structural pipeline; data collected during periods of head coil problems; data in which some of the FIX-ICA components were manually reclassified; participants that had any fMRI run in which more than 50% of TRs had greater than 0.25 mm motion framewise displacement; removal according to family relations (only unrelated participants were selected, and those with no genotype testing were excluded). This resulted in 352 participants, from which only the first half of participants (176) were included for computational tractability of our analyses. We include here a brief description of the data fMRI collection parameters: whole-brain echo-planar functional imaging acquisitions were acquired with a 32 channel head coil on a modified 3T Siemens Skyra MRI with TR = 720 ms, TE = 33.1 ms, flip angle = 52°, BW = 2290 Hz/Px, in-plane FOV = 208 × 180 mm, 72 slices, 2.0 mm isotropic voxels, with a multiband acceleration factor of 8. For our analysis, we only used one 14.4 minute run of resting-state data (1200 datapoints), and two 30 minute consecutive runs (60 min total) of task-state data (7 tasks with 24 conditions). Further task and resting-state data acquisition details can be found elsewhere (Barch et al., 2013; Smith et al., 2013).

In brief, the seven tasks consisted of an emotion cognition task (valence judgment, 2 conditions); gambling reward task (card guessing, 2 conditions); language processing task (2 conditions); motor task (tongue, finger, toe, 6 conditions); relational reasoning task (2 conditions); social interaction cognition task (2 conditions); and working memory task (0-back, 2-back, 8 conditions). Details about these task paradigms can be found in Barch et al. (2013).

The minimally-preprocessed HCP surface data (Glasser et al., 2013) were first parcellated into 360 cortical regions using the Glasser et al. (2016) atlas. Then, we applied the preprocessing steps detailed in Ito et al. (2020). Briefly, they include removing the first five datapoints of each run, demeaning and detrending the time series, and nuisance regression—based on Ciric et al. (2017)—with 64 parameters to control for the effects of motion and physiological artifacts, and their derivatives and quadratics. Global signal regression was not applied since its physiological basis and effects on functional connectivity inferences are still not fully understood (Aquino et al., 2020; Colenbier et al., 2020; Li et al., 2019; T. T. Liu et al., 2017; Murphy & Fox, 2017).

Task-evoked activations for each of the 360 regions and 24 conditions were estimated using a standard general linear model at the region level. The SPM software canonical hemodynamic response function (fil.ion.ucl.ac.uk/spm) was used for general linear model estimation given that all tasks involved block designs (Cole et al., 2021).

Analyses were run in the AmarelN cluster with the same specifications mentioned above. Data and code to reproduce our synthetic and empirical analyses are available at the project repository [available upon acceptance of the manuscript].

## Results

### Network recovery on simulated fMRI data

We began by simulating ground-truth functional causal networks to determine the accuracy of the different FC methods tested (**Figure 1**). A series of 200 random networks, each with 200 nodes, were simulated from a graphical causal model with more common causes and causal chains than colliders, and two-node and three-node cycles. Connectivity weights were sampled from a uniform distribution. For each causal network, we simulated fMRI time series with 1000 datapoints using a linear model. The accuracy of the FC methods was assessed in terms of how well they recovered ground-truth resting-state networks. We used precision and recall as measures of recovery accuracy.

We first report in **Figures 2A**-**C**, precision and recall for the recovery of the true network adjacency pattern in simulated data, for each of the FC methods tested. *Precision* is defined as the number of true-positive adjacencies (*tp*) divided by the sum of the number of true-positive and false-positive adjacencies (*fp*) (*precision* = *tp*/(*tp*+*fp*)). Precision values range from 0 to 1, and quantify the ability of each FC method to avert false-positive connections. A precision of 1 indicates that the method did not output any false positives, and a precision of 0 that it only output false-positive connections. *Recall* is defined as the number of true-positive adjacencies divided by the sum of the number of true-positive and false-negative adjacencies (*fn*) (*recall* = *tp*/(*tp*+*fn*)). Recall values also range from 0 to 1, and reflect the ability of each FC method to recover true connections. A recall of 1 indicates that the method inferred all the true connections, and a recall of 0 that it did not recover any of the true connections. Together, precision and recall yield a complementary view of each method’s capacity to recover the true network while avoiding false-positive connections. Results are reported in boxplots indicating median, and lower and upper quartiles for 200 simulated resting-state networks.

**Figure 2.**
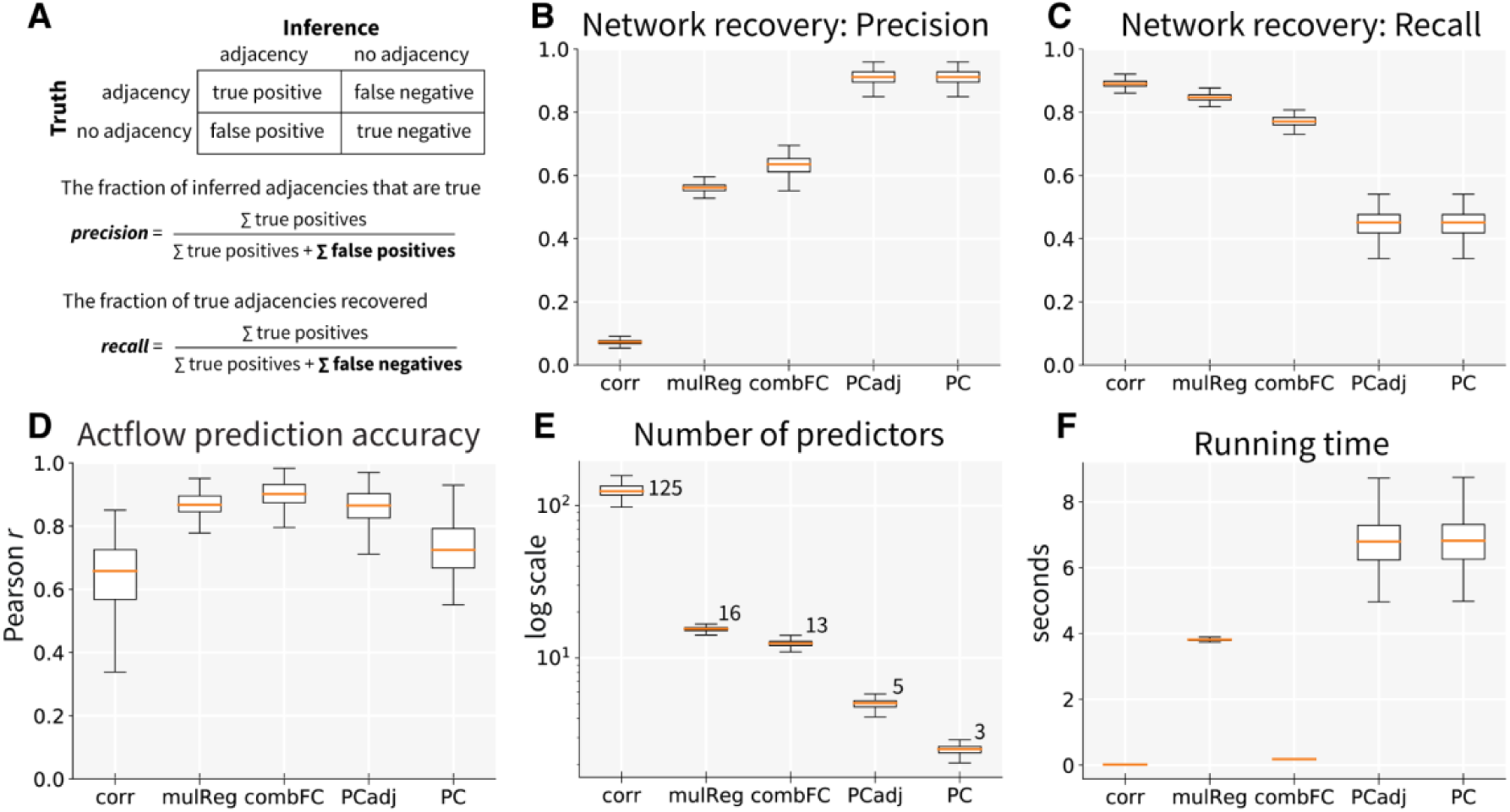
Recovery of functional connectivity networks and accuracy of activity flow prediction of task-evoked activations for simulated fMRI data. Boxplots show median, and lower and upper quartiles for 200 simulations. Correlation (corr), multiple regression (mulReg), combinedFC (combFC), PC-adjacencies (PCadj) and PC algorithm (PC). **(A)** Precision and recall formulas to measure goodness of network adjacency recovery. **(B)** Precision. **(C)** Recall. **(D)** Accuracy of actflow prediction of task-evoked activations, measured with Pearson correlation coefficient r and averaged across 200 regions. **(E)** Number of predictor regions averaged across 200 regions, plotted on a logarithmic scale for visualization, with actual median values next to each boxplot. For reference, the median in-degree (number of direct causal sources) in the true networks is 6.62. **(F)** Running time in seconds.

The preference to focus on reducing false-positive results (maximizing precision) or on reducing false-negative results (maximizing recall) critically depends on the research question being pursued. Here, we are mainly interested in measuring the performance of different FC methods in reducing the effect of deleterious spurious pathways and the subsequent impact on actflow predictions. With the primary goal of reducing the risk of false positive causal inferences, we sought to maximize precision first (reduce false-positive connections), followed by maximizing recall (reducing false negatives). Thus, the FC method with the highest precision would be preferred even with a moderately lower recall. Considering future practical applications involving prediction of causal interventions (e.g., brain stimulation effects), we are primarily interested in determining whether an effect would occur when it is predicted. This led us to prioritize avoiding cases where a false positive indicates a causal effect would occur when in fact it would not once an intervention is applied (e.g., in a randomized controlled trial). While this led us to avoid false positives over false negatives, it will be important for future work to also reduce false negatives, given the potential for interventions to have undesirable side effects that are predicted only in the absence of false negatives.

As stated above, we recognize that for some research questions it may be more important to minimize false negatives than false positives. For these cases we suggest two practical alternatives. The first one is to increase the α threshold of the PC algorithm, which may result in a larger number of connections thus increasing the recall (reduction of false-negative connections) at the cost of losing precision (increase of false-positive connections). The second alternative is to use combinedFC, which in simulations showed better recall but lower precision than the PC algorithm (**Figures 2B**-**C**). In both alternatives, researchers should consider how the reduction in false negatives may improve the mechanistic interpretation of their models (for example, by including a more complete description of the true source connections of a target region), but also the cost of increasing false positive inferences (for example, the higher risk of a failed intervention informed by false-positive connections).

Considering the theoretical properties of each FC method to avoid false-positive connections from confounders, chains and conditioned-on colliders, we expect the PC algorithm to achieve the highest precision (less false-positive connections) for network recovery, followed by combinedFC, multiple regression and correlation, in that order. Regarding recall, it is not clear which of the FC methods is expected to recover the highest number of true connections.

**Figure 2B** shows that, as expected, PCadj and PC dominated over the rest of the methods with a median precision of 0.91, indicating that 91% of its inferred adjacencies are true-positive connections and 9% are false-positive connections. Note that PCadj and PC have the exact same precision since PCadj is PC without the orientation information, and here precision is calculated only for adjacencies. CombinedFC and multiple regression were next, with precisions of 0.64 and 0.56 respectively. The lowest scoring method was correlation, with a precision of 0.07, implying that 7% of its inferred adjacencies are true positives and 93% are false positives.

**Figure 2C** shows that the results for recall go in the opposite direction. Correlation had the highest median recall, 0.89, indicating that 89% of the total true connections were recovered. Multiple regression and combinedFC follow closely with 85% and 77% respectively, while PCadj and PC had the lowest recall, with 45% of the true connections recovered.

From these methods, only the PC algorithm has inference rules to orient connections (**Box 1 (2)**). To measure the orientation accuracy of PC, meaning if the causal direction of an adjacency was correctly inferred, we computed the proportion of correct orientations out of the total number of correctly inferred adjacencies (true-positive adjacencies). Notably, PC showed a median orientation accuracy of 0.83 across the 200 simulations, implying that 83% of the true-positive adjacencies inferred were oriented in its correct causal direction.

These first results confirm that FC methods that make use of more conditional independence information and causal principles, such as combinedFC and PC, can recover with higher precision (lower number of false-positive connections) simulated resting-state fMRI networks. This suggests that these FC methods will result in activity flow models with better prediction accuracy, since the prediction of the task-evoked activations will not contain effects from spurious pathways.

### Actflow prediction accuracy on fMRI synthetic networks

We extended our ground-truth models to determine if FC methods that better control the effect of false-positive connections, lead to more accurate predictions of task-evoked activity. Task-state networks were simulated by taking the previous 200 resting-state simulated networks and applying minor deviations to the original connectivity weights. For each iteration, the task was modeled as an exogenous source node, randomly connected to a number of network nodes. Task fMRI time series with 1000 datapoints were simulated for each network. We regressed each task-state network node on the exogenous task and considered the regression coefficients as the to-be-predicted (actual) task-evoked activations.

In **Figure 2D** we report the prediction accuracy of the actflow models for each of the FC methods tested. Following Cole et al. (2016), we measured prediction accuracy with the Pearson correlation coefficient *r* between the vector of predicted activations for the 200 simulated regions and the vector of actual activations. Measured in this way, the prediction accuracy *r* summarizes the actflow model performance across the whole network. Boxplots show median, and lower and upper quartiles for 200 simulations.

Previous fMRI empirical results (Cole et al., 2016) showed that functional networks estimated with multiple regression produced more accurate actflow predictions compared to networks estimated with correlation. In addition, our simulation results (**Figures 2A** and **2B**) show that multiple regression-based networks have a higher number of false negatives (lower recall) but a lower number of false positives (higher precision) than correlation-based networks. Together, these observations suggest that actflow prediction accuracy may be more improved by reducing false-positive functional connections than by reducing false-negative connections. Thus, we expect that PCadj and PC—the methods with the best network recovery precision—will have the higher prediction accuracy, followed by combinedFC, multiple regression and correlation, in that order.

CombinedFC had the best median prediction accuracy of all the methods (*r* = 0.90), probably due to a good balance for inferring real pathways and avoiding spurious pathways. It was followed closely by PCadj (*r* = 0.87) and multiple regression (*r* = 0.87). PC had a lower prediction accuracy compared to these methods (*r* = 0.72). Note that when the actflow model is parameterized with the PC-based oriented network, it only uses the estimated direct sources to predict the activity of each held-out region (biologically-constrained mechanistic actflow). In contrast, when actflow uses unoriented networks, derived from multiple regression, PCadj or combinedFC for example, it considers all the adjacent regions of a held-out region to predict its activity. In causal terms, this implies that with unoriented networks, the held-out region activity prediction leverages information from both direct sources and direct effects, achieving a better prediction than when only source information is used. It is for this reason that the PC prediction accuracy was lower than PCadj, multiple regression and combinedFC prediction accuracies. Critically, the higher network recovery precision for the PC-based models (**Figure 2B**) means that (despite having lower prediction accuracy) they provide increasing mechanistic interpretability of the actflow-generated activity predictions.

As expected, actflow models based on correlation networks showed the lowest accuracy of all (*r* = 0.66). Despite the low number of false negatives (high recall), these models have a high number of false-positive connections (low precision), which create spurious pathways through which task activity is incorrectly accounted for.

Note that a vector of predictions and a vector of actual activations can have a high Pearson *r*, even if their values are in a completely different scale (e.g., all values multiplied by 2). If we are interested in assessing the deviation from the actual values, we can use the coefficient of determination *R*^2^ = 1−(Σ*_i_*(*A_i_*−*Â_i_*)^2^/Σ*_i_*(*A_i_*−*Ā*)^2^), where *A_i_* are the actual activations, *Â_i_* the predicted activations, and *Ā* the mean of the actual activations. *R*^2^ measures the proportion of the variance of the actual activations that is explained by the prediction model. It ranges from 1 (perfect prediction) to minus infinity (prediction deviations can be arbitrarily large), with a value of zero when the predictions are equal to the mean of the actual activations (*Â_i_* = *Ā*). For our simulations, the median actflow prediction *R*^2^ for the FC methods followed the same ordering as the Pearson *r*: combinedFC (*R*^2^ = 0.80), PCadj (*R*^2^ = 0.74), multiple regression (*R*^2^ = 0.68), PC (*R*^2^ = 0.52) and correlation (*R*^2^ = −164.68). The high negative *R*^2^ of correlation-based models indicates that predicted values strongly deviate from actual activation values, confirming the detrimental effect of spurious network pathways on the actflow prediction model.

We evaluated the complexity of actflow models with the number of regions used to predict the task-evoked activation of held-out regions. In practice, as the complexity of a model increases, its interpretability decreases, making it more difficult to parse out the network processes underlying the generation of task-related activations. **Figure 2E** shows the number of predictors for each held-out region, averaged across the 200 regions. Notably, actflow models based on PC directed networks achieved high accuracy (**Figure 2D**), with the lowest model complexity (median of 3 predictors, since only directed sources were used to predict activations). These results evidence that PC can successfully recover functional directed connections that have high predictive power. This is further confirmed with the results of PCadj-based actflow models, which also had a relatively low number of predictors (median of 5), and an accuracy as high as combinedFC and multiple regression, both with an order of magnitude more predictors (median of 13 and 16 respectively). Correlation-based actflow models reported the highest complexity (median of 125 predictors) and the lowest accuracy to predict task-evoked activations.

Finally, **Figure 2F** shows that all FC methods have efficient running times which do not exceed the tens of seconds. The PC algorithm had the longest median running time (7 sec), which is very efficient considering the large number of conditional associations it has to compute due to the number of regions and the complexity of the connectivity patterns in the true networks. Surprisingly, multiple regression had a relatively long median running time (4 sec). Analysis of our code revealed that this unexpected running time was caused by the significance test for the multiple regression coefficients. Not computing the significance test considerably reduces the running time. Running times depend on the hardware used for the analysis, but we expect that the reported ordering of the methods will replicate in any machine.

Our results on simulated fMRI data show that PC and combinedFC can be used to build activity flow models with high prediction accuracy, low complexity (number of predictors) and efficient running times. Thanks to their use of statistical conditional independence information and causal principles to control for the effects of confounders, causal chains and conditioned-on colliders, and if orientation rules are provided, such as with PC, these methods can provide plausible mechanistic hypotheses regarding the generation of task-evoked activations.

### Actflow prediction accuracy on fMRI empirical networks

Our theoretical considerations about the continuum of FC methods (**Figure 1**) and the comparative performance observed in simulations, prompted us to hypothesize that this performance will translate, up to a degree, to an empirical domain. Here, we used fMRI data to assess the performance of actflow predictive models under a complex empirical setup—for which the ground-truth networks are not known—comprising 360 cortical regions, 24 task conditions across various cognitive domains and 176 different participants. We compared the average prediction accuracy of the different FC methods across all conditions.

Prediction accuracy was measured as the Pearson correlation coefficient *r* between the vector of predicted activations for all 360 cortical regions of Glasser et al. (2016) and the vector of actual activations. We computed the prediction accuracy individually for each of the 24 HCP task conditions and reported the average across conditions. Measured in this way, the prediction accuracy *r* summarizes the actflow model performance across the whole brain and the full set of task conditions. **Figure 3A** reports actflow prediction accuracy boxplots with median, and lower and upper quartiles across the 176 participants, for each FC method. PCadj attained the highest median prediction accuracy (*r* = 0.82), followed by combinedFC (*r* = 0.77) and PC (*r* = 0.74). Multiple regression (*r* = 0.60) and correlation (*r* = 0.57) showed lower accuracies. As in the simulation results, we include median coefficient of determination *R*^2^ as a complementary measure of prediction accuracy: PCadj (*R*^2^ = 0.67), combinedFC (*R*^2^ = 0.59), PC (*R*^2^ = 0.53), multiple regression (*R*^2^ = 0.33) and correlation (*R*^2^ = −710). As remarked in the simulation results, the high negative *R*^2^ of the correlation-based models reflects large differences between the predicted and the actual activation values, which are likely the result of spurious network pathways. As we hypothesized, these empirical results are consistent with the simulations, in the sense that actflow models parameterized with FC methods that use more statistical conditional independence information and causal principles, such as PC and combinedFC, can better predict task-evoked activity.

**Figure 3.**
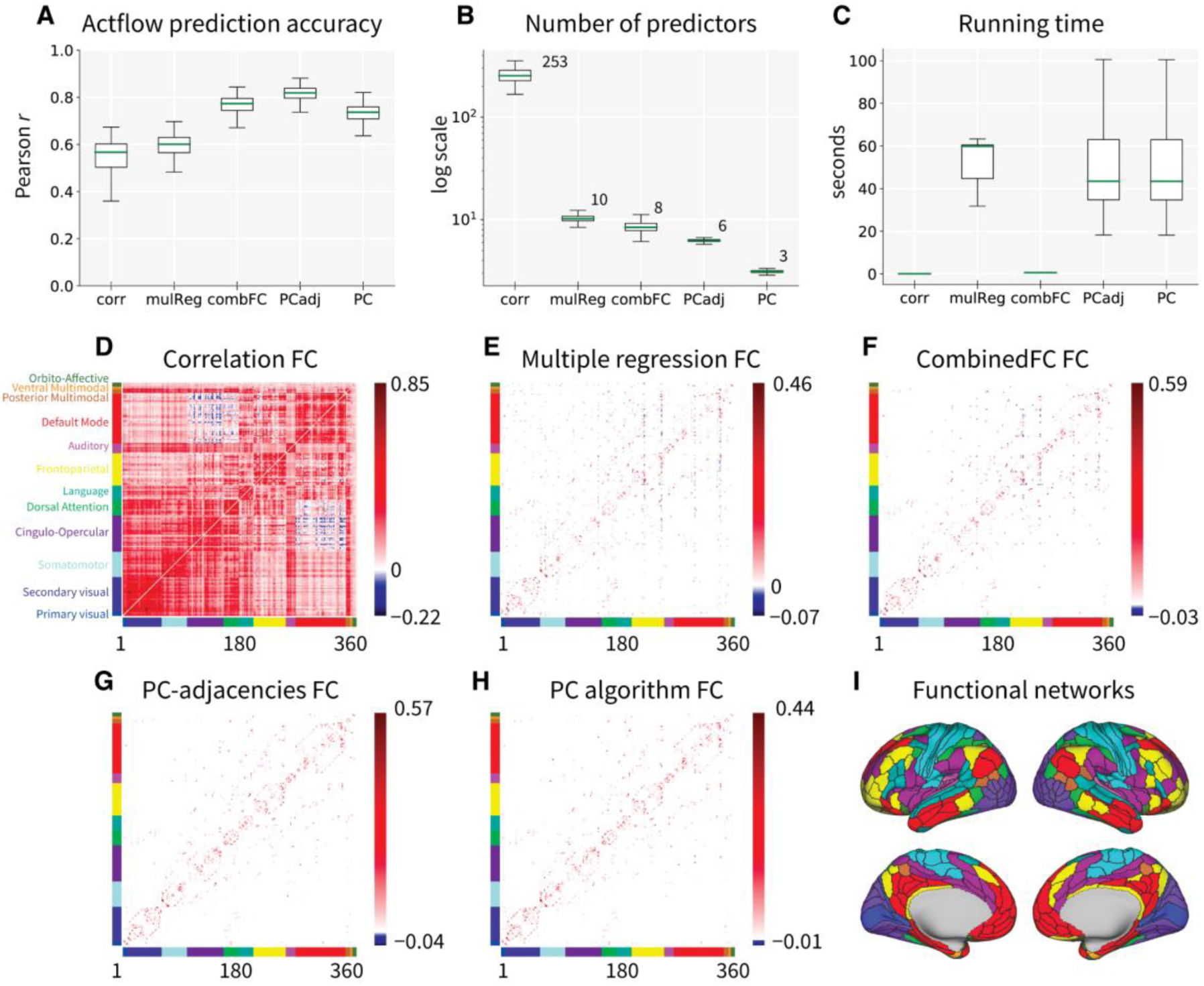
Accuracy of activity flow prediction of task-evoked activations based on functional connectivity networks estimated from empirical fMRI data. Correlation (corr), multiple regression (mulReg), combinedFC (combFC), PC-adjacencies (PCadj) and PC algorithm (PC). Boxplots show median, and lower and upper quartiles for the 176 participants. **(A)** Prediction accuracy for activity flow models parameterized with each of the FC methods tested. Accuracy was measured as the Pearson correlation coefficient r between the vector of predicted activations for all 360 cortical regions of Glasser et al. (2016) and the vector of actual activations, averaged across the 24 HCP task conditions. **(B)** Number of predictor regions averaged across 360 regions, plotted on a logarithmic scale for better visualization, with actual values next to each boxplot. **(C)** Running time in seconds for network estimation. **(D)** Correlation-based functional connectivity (FC) network, averaged across 176 participants. The 360 cortical regions were organized in 12 functional networks from Ji et al. (2019). FC average network for **(E)** multiple regression, **(F)** combinedFC, **(G)** PC-adjacencies and **(H)** PC algorithm. **(I)** Cortical surface map of the 12 functional networks partition used in panels D-H (available at https://github.com/ColeLab/ColeAnticevicNetPartition). For visualization, in all FC networks (panels D-H), values between -/+ 0.005 were set to zero.

The complexity of the actflow models (measured as the number of predictors) (**Figure 3B**) followed the same order observed in simulations. PC had the lowest model complexity (median of 3 predictor source regions), followed by PCadj (median of 6), combinedFC (median of 8), multiple regression (median of 10) and correlation (median of 253). These results also reflect the different connectivity densities of the estimated FC networks. Correlation produced highly dense networks (**Figure 3D**), resulting in actflow models with high complexity (large number of predictors) but low prediction accuracy (**Figure 3A**), whereas PC inferred sparser networks (**Figures 3G**-**H**), that produced actflow models with lower complexity and high prediction accuracy.

To confirm that the good performance of sparser PCadj-based actflow models comes from the appropriate control of confounders, causal chains and condition-on colliders, and not just from their sparser inferred networks, we report actflow prediction accuracy across undirected FC methods after matching their connectivity density (and median number of predictors) to that of PCadj networks. To do this, we thresholded networks by selecting the connections with the strongest absolute value until the PCadj density was matched. As above, we use median Pearson correlation and coefficient of determination to measure accuracy, and compare these values against the previously observed accuracies from Figure 3. We observed a significant reduction in accuracy for correlation (*r* = 0.48 vs. 0.57) (but increase in *R*^2^ = -33 vs. -710, probably due to a reduction in predicted activation values); a significant reduction for multiple regression (*r* = 0.53 vs. 0.60, *R*^2^ = 0.23 vs. 0.33); and no significant change for combinedFC (*r* = 0.77 vs. 0.77, *R*^2^ = 0.59 vs. 0.59) (*p* < 0.0001 for all significant differences, for a two-sample *t*-test one-sided, *n* = 176). All of these methods retained lower prediction accuracy than PCadj (*r* = 0.82, *R*^2^ = 0.67), despite matching PCadj network density. These results confirm that PCadj-based actflow models’ performance comes from the effective leverage of statistical conditional independence information and causal principles, and cannot be replicated in the other methods by forcing network sparsity via an arbitrary connection-strength-based thresholding approach.

Lastly, **Figure 3C** reports running times for the inference of FC networks from empirical fMRI time series. We observed efficient running times not exceeding tens of seconds. For these data, multiple regression showed the longest median running time (44 seconds), followed by PC (33 seconds). As mentioned above, the relatively long running time of multiple regression comes mainly from the computation of the significance test for the regression coefficients.

Together, these results confirm empirically that methods using statistical conditional independence information and causal principles can estimate undirected and directed FC networks useful to build activity flow models with low complexity and high prediction accuracy. Actflow models using undirected FC networks can provide excellent predictions (e.g., combinedFC and PCadj), but cannot tell if the task activity is flowing into, from, or into and from (as in a feedback) the held-out region. In contrast, PC-based directed networks, despite reduction in the prediction accuracy, provide biologically-constrained mechanistic models in which the flow of task activity can be traced from source regions directly into target regions.

### Actflow predictions across brain regions and task conditions

Our results so far have confirmed in simulations and empirically (**Figure 3A**) the benefits in prediction and mechanistic interpretation from FC methods that incorporate statistical conditional independence information and causal principles. Notably, we showed that directed FC networks from the PC algorithm can be used to build directed actflow models (**Figure 1H**) with high prediction accuracy, low complexity and tractable mechanistic interpretation. To further illustrate the benefits of PC-based directed actflow models, we compare their performance against the field-standard Pearson correlation FC, the method at the extreme of our continuum, only based on statistical pairwise associations. This analysis is essentially focused on measuring the accuracy in predicting a whole-brain pattern of regional activations for each task condition (node-wise accuracy) (Cole et al., 2021).

The actflow accuracies reported in **Figure 3A** summarize on the Pearson *r*, general information from all 360 cortical regions and 24 task conditions. Here, we unpacked this information by showing in matrix form the actual task activations for all regions (rows) and task conditions (columns), median across the 176 participants (**Figure 4B**), flanked by the predicted activations of the field-standard Pearson correlation-based models (**Figure 4A**) and by the predictions of the PC-based mechanistic actflow models (**Figure 4C**). These matrices show with greater detail how PC-based predictions across regions and all conditions had a more similar pattern to the actual activations, relative to correlation-based predictions (*r* = 0.74 > *r* = 0.57, median across participants). Importantly, we also observed that the PC-based predicted values were in approximately the same range as the actual activations (compare colorbars in **Figures 4B** and **4C**), while correlation-based predicted activations were often hundreds of times off of the actual values (see **Figure 4A** colorbar). As mentioned above, these deviations can be quantitatively assessed with the coefficient of determination, *R*^2^ = 0.53 vs. *R*^2^ = −710, for PC-based models and correlation-based models correspondingly. The high negative *R*^2^ reflects the observed strong deviations in the correlation-based actflow predicted values.

**Figure 4.**
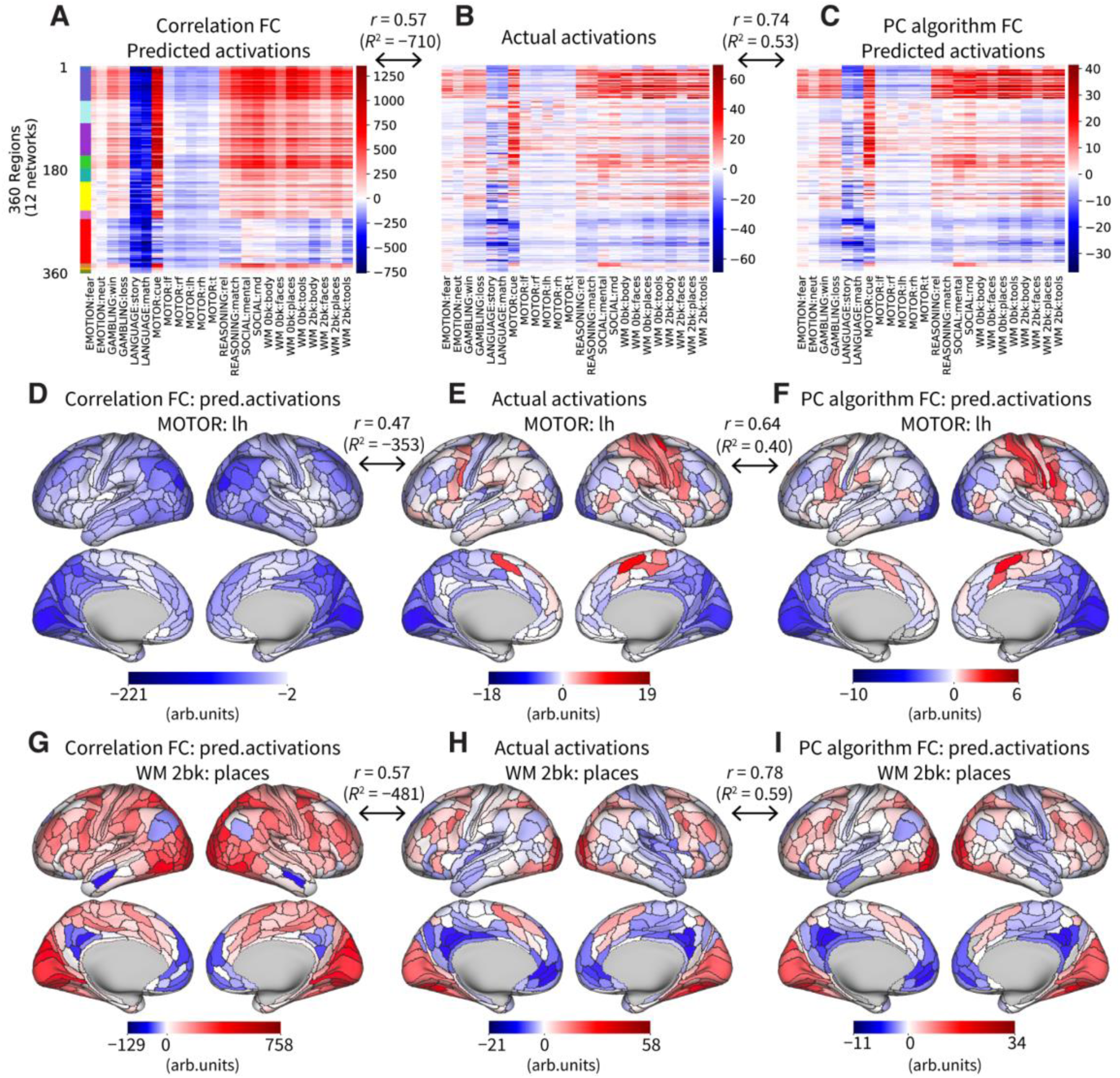
Activity flow prediction of task-evoked activations for empirical fMRI data, across cortical regions and task conditions. **(A)** Actflow predicted activations based on correlation FC, for 360 regions organized in 12 networks (rows) (Glasser et al., 2016; Ji et al., 2019), and 24 HCP task conditions (columns), median across 176 participants. The median results from first computing the Pearson *r* (or coefficient of determination *R*^2^) prediction accuracy node-wise (360 regions) for each of the 24 conditions, then averaging across conditions and finally obtaining the median across participants. **(B)** Actual task-evoked activations. **(C)** Actflow predicted activations based on PC algorithm FC. **(D)** Correlation-based actflow predictions for the motor task: left hand condition, projected into a brain surface map with 360 Glasser regions, median across 176 participants. The prediction accuracy is the Pearson *r* (or coefficient of determination *R*^2^) between the vector of node-wise (360 regions) predicted activations for this condition and the vector of actual activations, median across participants. **(E)** Actual activations for the motor task: left hand condition. **(F)** PC-based actflow predictions for the motor task: left hand condition, and median prediction accuracy. **(G-I)** Same results as panels D-F but for working memory task: 2-back: places condition.

For each of the 24 task conditions taken individually, we confirmed that PC-based actflow models attained a significantly higher node-wise prediction accuracy than correlation-based models (for both Pearson *r* and *R*^2^, *p* < 0.01 corrected for multiple comparisons with the nonparametric test of Nichols & Holmes (2002), for 176 participants).

To further illustrate this result, we performed a more targeted analysis that focuses on one task condition (one column in the **Figures 4A**-**C** matrices) and examines the pattern of actflow predicted activations across the 360 brain regions (node-wise accuracy). We first show this analysis for the motor task: left hand condition. The PC-based actflow models (**Figure 4F**) recovered the whole-brain actual activation pattern (**Figure 4E**) better than the correlation-based models (**Figure 4D**) (*r* = 0.64 > *r* = 0.47; and *R*^2^ = 0.40 vs. *R*^2^ = −353, median across participants). More importantly, the PC-based models correctly recovered the functionally-relevant pattern of positive activations in the right hemisphere somatomotor network, known to be engaged during task-induced left hand movements. In contrast, correlation-based models predicted negative and inflated activation values across the entire cortex.

We repeated the node-wise accuracy analysis, this time for the working memory task: 2-back: places condition. In this case, we also confirmed the superior performance of PC-based actflow models (**Figure 4I**) compared to correlation-based models (**Figure 4G**), both in the prediction of the whole-brain activation pattern and in the range of predicted values (*r* = 0.78 > *r* = 0.57; and *R*^2^ = 0.59 vs. *R*^2^ = −481, median across participants). Note how the predictions of correlation-based models are massively biased towards positive and negative values (**Figure 4G**, colorbar), reflecting again the presence of inferred spurious pathways through which task activity is incorrectly summed to the actflow computation.

These results confirm that mechanistic actflow models based on FC methods, that control for indirect and spurious pathways, and provide causal source information, such as PC, better predict whole-brain activation patterns and actual values, for each of the 24 task conditions tested here. In contrast, correlation-based FC produced densely connected actflow models in which every region’s connectivity profile probably conflated direct, indirect and spurious pathways (see **Figure 3D** FC matrix) that incorrectly biased the actflow predicted activations.

### Region-wise activity flow prediction across task conditions

The previous analysis compared the accuracy of PC- and correlation-based actflow models in predicting whole-brain activation patterns for individual task conditions (node-wise accuracy, e.g., **Figure 4F**). Here, in contrast, we want to measure the accuracy of PC-based directed actflow models and correlation-based models for predicting activations across the 24 task conditions for each individual brain region (condition-wise accuracy) (Cole et al., 2021). This analysis allows us to highlight brain areas for which the two methods differ significantly.

We consider the matrices in **Figures 4A**-**C**, choosing one row (region) and computing the Pearson *r* value between the vector of 24 column (conditions) actual activations and the vector of 24 column actflow-predicted activations. The prediction accuracy *r* value for each region is then projected to a brain map. **Figures 5A**-**B** show the condition-wise accuracy of correlation-based and PC-based actflow models for each region (median across 176 participants).

**Figure 5.**
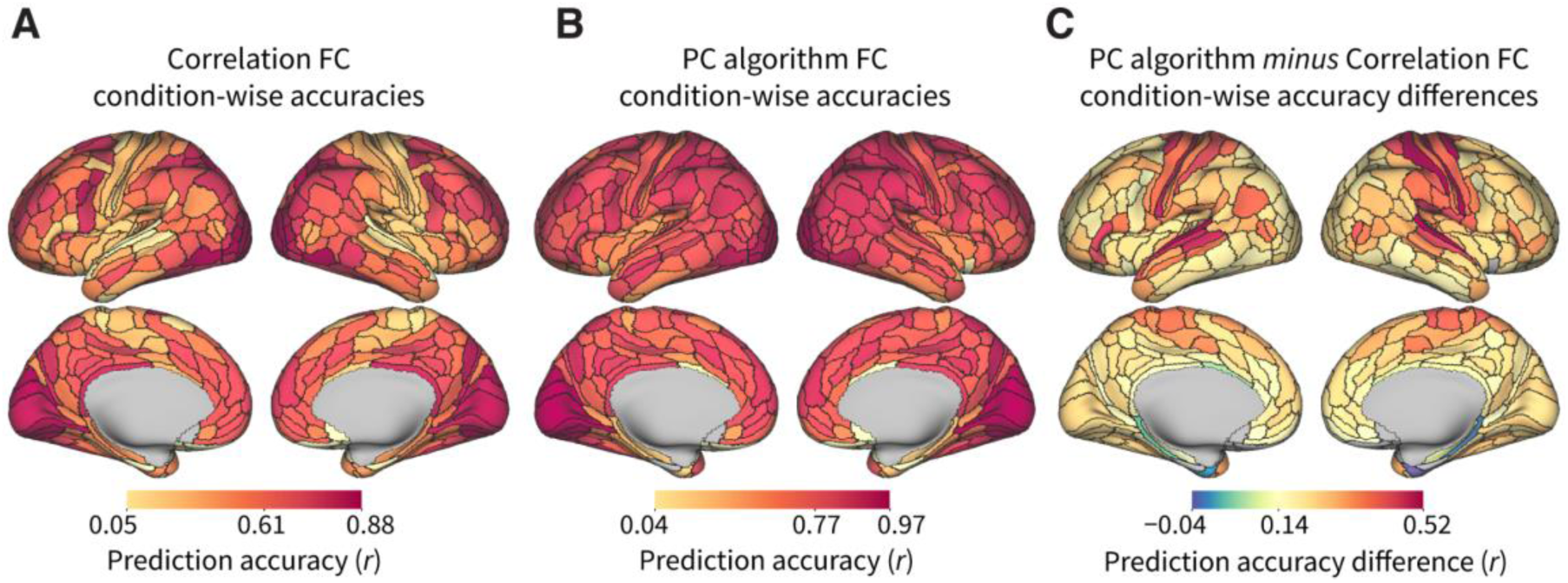
Condition-wise accuracy of activity flow predictions for 360 cortical regions. Condition-wise actflow accuracy is the prediction accuracy for each region across the 24 task conditions of the HCP data. Brain surface maps show median across 176 participants. Colorbars indicate minimum, mean and maximum prediction accuracy (r) across the 360 Glasser regions. **(A)** Actflow condition-wise accuracies using the field-standard Pearson correlation FC. **(B)** PC algorithm-based actflow condition-wise accuracies. **(C)** PC algorithm-based minus correlation-based actflow condition-wise accuracies. Positive values indicate that PC algorithm-based actflow models predict better than correlation-based models, negative values indicate the opposite.

To highlight differences in prediction accuracy across the methods, **Figure 5C** shows the PC-based condition-wise accuracy minus the correlation-based accuracy for each brain region. For 82% of the 360 regions, PC-based actflow models attained a significantly higher condition-wise prediction accuracy than correlation-based models (Pearson *r*, *p* < 0.01 corrected for multiple comparisons with the nonparametric test of Nichols & Holmes (2002), for 176 participants. 99% of the 360 regions for *R*^2^). The brain map of **Figure 5C** shows that accuracy differences are relatively larger in the language and somatomotor networks. This condition-wise result is consistent and complementary to the previous node-wise accuracy analysis of the motor and working memory tasks (**Figures 4D**-**I**).

These results extend our previous observations by confirming that PC-based directed actflow models (with lower complexity and valid mechanistic interpretation) can better predict, for a large majority of cortical regions, the task-evoked activations across a diverse set of cognitive conditions.

### Demonstrating how directed activity flow models can improve mechanistic insight into the generation of cognitive functions: DLPFC working memory condition selectivity

Results thus far demonstrate that parameterizing activity flow models with FC methods that effectively exploit conditional independence information and causal principles improves our ability to accurately simulate the generation of task-evoked activations. This was shown in previous sections via summaries of results across all 360 regions and 24 task conditions of our empirical dataset. Here, to illustrate how actflow models can be used in practice to advance mechanistic explanations of cognitive function, we focus on one particular brain region engaged in a single cognitive task manipulation. Specifically, we applied an actflow model to understand how the flow of activity from distal brain regions may give rise to an established cognitive activation effect (2-back vs. 0-back working memory task conditions) in one region of the right dorsolateral prefrontal cortex (DLPFC). We focused on a cognitive contrast given the enhanced causal experimental control inherent in such a contrast (controlling for, e.g., stimulus perception, task timing, general task engagement). To further illustrate the impact that the choice of FC method can have on actflow-based explanations, we again compare results for correlation-based (the field standard and where no conditional independence information or causal principles are used) and PC-based actflow models (where the largest amount of conditional independence information and causal principles are used).

First, activations for the 4 conditions for 2-back (body, face, place and tool stimuli) (see **Figures 4A**-**C**) were averaged to form one 2-back activation for each of the 360 Glasser regions. The same procedure was applied for the 0-back conditions. Then, activation contrasts (2-back minus 0-back) were computed for each subject, and regions with higher 2-back than 0-back activation on average across subjects were localized (**Figure 6A**, red regions).

**Figure 6.**
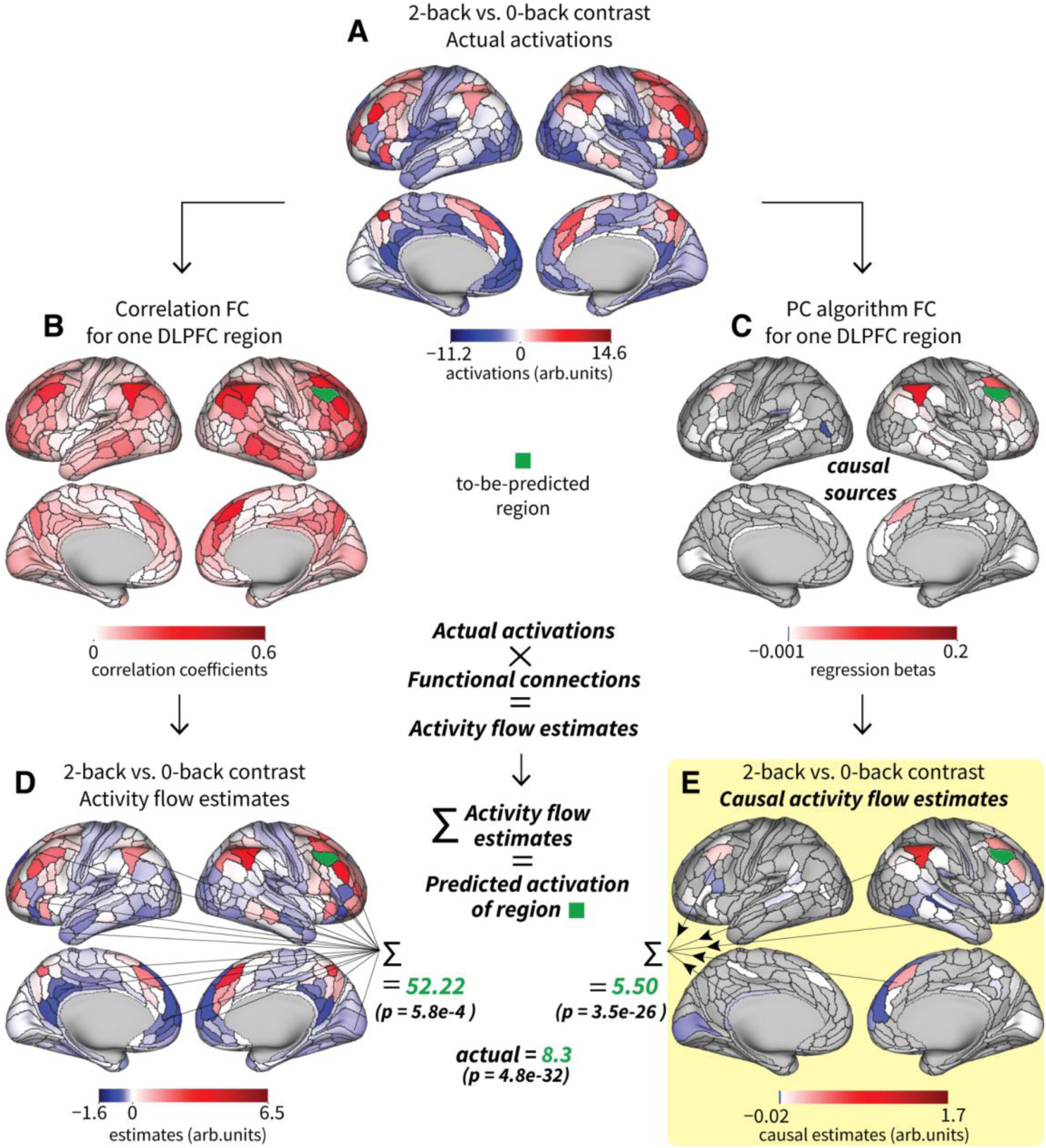
Directed activity flow models provide mechanistic insight into the generation of n-back cognitive responses in DLPFC. **(A)** 2-back vs. 0-back actual activation contrast, for each of the 360 Glasser cortical regions. **(B)** Correlation FC for one to-be-predicted region in the right hemisphere dorsolateral prefrontal cortex (green region, right DLPFC Area 8C), averaged across 176 participants. **(C)** PC algorithm directed FC from putative causal sources to the to-be-predicted right DLPFC Area 8C (green region). **(D)** Correlation-based activity flow estimates, resulting from multiplying each region’s actual activation contrast (panel A) by its correlation functional connection (panel B), averaged across 176 participants. The region-wise sum of the activity flow estimates is the correlation-based actflow contrast prediction for the right DLPFC Area 8C; average predicted value across 176 participants, 52.22, *p*-value = 5.8e-4 for a two-sided *t*-test. **(E)** PC algorithm-based causal activity flow estimates, resulting from multiplying contrasts (panel A) by its PC directed functional connection (panel C). Average predicted value across 176 participants, 5.50, *p*-value = 3.5e-26 for a two-sided *t*-test. The actual activation contrast for the right DLPFC Area 8C is included for comparison, average across participants 8.3, *p*-value = 4.8e-32 for a two-sided *t*-test.

Brodmann’s Area 8 in the posterior section of the DLPFC has been extensively reported to have an important role in the maintenance of stimulus information during working memory tasks (Carlson et al., 1998; Constantinidis et al., 2001; Courtney et al., 1998; Petrides, 2000; Rowe et al., 2000; Rowe & Passingham, 2001; Wager & Smith, 2003). In the Glasser et al. (2016) parcellation, DLPFC Area 8 is subdivided into areas 8C, 8Av, 8Ad and 8BL, from which the right hemisphere Area 8C showed the highest positive activation in the 2-back vs. 0-back contrast—both in the analysis conducted here and the one in Glasser et al. (2016). The right DLPFC Area 8C therefore has a prominent and established role in working memory function, and this motivated us to choose it as our to-be-predicted target (**Figure 6C**, green region).

We built an actflow model for right DLPFC Area 8C using the actual activation contrasts for the rest of the regions (**Figure 6A**) and its estimated resting-state functional connections (**Figures 6B** and **C**). We obtained a brain map of activity flow contrast estimates by multiplying each region’s actual activation contrast by its corresponding connection to the to-be-predicted target (**Figures 6D** and **E**). Positive values indicate incoming contributions that would increase activity in right Area 8C, while negative values indicate incoming contributions that would decrease activity in this region. Finally, we summed the activity flow estimates to simulate the generation of the task-evoked activation contrast for the chosen DLPFC region.

Correlation-based actflow models used, on average, a considerably high number of regions in the left and right hemispheres (**Figure 6B**) to predict the activation contrast of the target right DLPFC Area 8C. These densely connected models predicted an activation contrast one order of magnitude larger than the actual contrast (52.22 vs. 8.3, average across 176 participants) (**Figure 6D**). In contrast, the PC algorithm inferred, on average, directed and sparser connectivity patterns for the DLPFC Area 8C (**Figure 6C**). These sparser patterns resulted in an actflow prediction closer to the actual contrast value (5.45 vs. 8.3, average across 176 participants) (**Figure 6E**). In line with these observations, the mean absolute error (MAE) for correlation-based actflow models was significantly larger than the MAE for PC-based models (*p* = 7.4e-31 for a Wilcoxon signed-rank non-parametric test, *n* = 176).

In addition, we identified 18 regions distributed across the default mode network (3 regions) and the frontoparietal network (15 regions), whose group-average absolute-value (to consider positive and negative estimates) actflow estimates are at the 95th percentile of the distribution, suggesting a relevant contribution of these afferent regions in the network-supported generation of working memory effects in Area 8C of the right DLPFC (**Table 2**).

**Table 2.** Relevant contributing regions to the PC-based actflow DLPFC prediction. 18 regions (out of 360) with the strongest contributions (activity flow estimates) to predict right DLPFC Area 8C 2-back vs. 0-back working memory contrast. Group-average absolute-value activity flow estimates are at the 95th percentile of the distribution. Region name and location from (Glasser et al., 2016), network assignment from (Ji et al., 2019).

| Region | Cortical location | Network | Activity Flow Estimate |
| --- | --- | --- | --- |
| Area PFm Complex | Inferior Parietal Cortex | Frontoparietal (FPN) | +1.682 |
| Area 8Av | Dorsolateral Prefrontal Cortex | Default Mode (DMN) | +0.791 |
| Area 8BM | Anterior Cingulate and Medial Prefrontal Cortex | FPN | +0.712 |
| Area posterior 9-46v | Dorsolateral Prefrontal Cortex | FPN | +0.567 |
| Area 8C | Dorsolateral Prefrontal Cortex | FPN | +0.343 |
| Inferior 6-8 Transitional Area | Dorsolateral Prefrontal Cortex | FPN | +0.220 |
| Area anterior 47r | Orbital and Polar Frontal Cortex | FPN | +0.135 |
| Area IFJp | Inferior Frontal Cortex | FPN | +0.087 |
| Area posterior 10p | Orbital and Polar Frontal Cortex | FPN | +0.086 |
| Area PGs | Inferior Parietal Cortex | DMN | +0.083 |
| Area 44 | Inferior Frontal Cortex | FPN | +0.083 |
| Area 9 Posterior | Dorsolateral Prefrontal Cortex | DMN | +0.076 |
| Area IntraParietal 1 | Inferior Parietal Cortex | FPN | +0.075 |
| Area IFSp | Inferior Frontal Cortex | FPN | +0.068 |
| Anterior Ventral Insular Area | Insular and Frontal Opercular Cortex | FPN | +0.060 |
| Superior 6-8 Transitional Area | Dorsolateral Prefrontal Cortex | FPN | +0.058 |
| Area anterior 9-46v | Dorsolateral Prefrontal Cortex | FPN | +0.051 |
| Area posterior 47r | Inferior Frontal Cortex | FPN | +0.051 |

Activity flow models also allow us to quantify the percentage of inter-subject variance in task-related activity that network distributed afferent processes can explain. We quantify this for Area 8C of the right DLPFC, using the *R*^2^ between the PC-based actflow model predictions and the actual n-back-related contrasts across individuals. Data-driven causally-informed actflow models including distributed activity from regions in default mode and frontoparietal networks were able to explain 47% of variance across participants (*R*^2^ = 0.47, *n* = 176). This value was much lower (*R*^2^ = -693.47) for correlation-based actflow models. This result supports the conclusion that a substantial portion of right DLPFC Area 8C activation variance can be explained in terms of distributed network processes across the default mode and frontoparietal networks, and that PC-based connectivity estimates could be used to more accurately model that variance relative to standard correlation-based connectivity estimates.

The information provided by the directed activity flow models of DLPFC during a working memory task (**Figure 6** and **Table 2**) can be further used to expand our mechanistic explanation of the distributed processes supporting working memory. For example, following Mill et al. (2022), and Hearne et al. (2021), we can use actflow with resting-state FC between regions in **Table 2** and DLPFC in a clinical population with working memory deficits to predict unhealthy task-evoked working memory activations. These predicted activations can be further used in a decoding model of behavior to explore the causal relations between FC changes and working memory deficits. Another possibility is to follow Ito et al. (2022), using resting-state FC patterns as a part of a multi-step empirical neural network model that starts in regions encoding working memory task stimuli, then follows to intermediate processing of task-relevant information (such as the one occurring between regions in **Table 2** and DLPFC) and finishes in motor regions where the response demanded by the task is decoded, usually button presses. A third possibility is to use non-invasive approaches such as transcranial electrical stimulation to alter the connectivity (Reinhart, 2017) between regions in **Table 2** and DLPFC and measure the effect of FC alterations on the activation patterns and behavioral response during the working memory task.

## Discussion

We used simulations and empirical fMRI to test the hypothesis that FC methods that use statistical conditional independence information and statistical causal principles, can accurately recover causal properties of functional brain networks that can then be used to construct activity flow models—empirically-constrained network simulations. These empirically-constrained simulations provide plausible distributed network accounts of the generation of cognitive brain activations. Importantly, we illustrated the explanatory potential of activity flow models with a PC-based actflow model of right DLPFC (Area 8C), which provides a unique distributed account of an established cognitive effect observed during increased working memory load (a statistically significant 2-back vs. 0-back contrast) (Sreenivasan et al., 2014).

Specifically, our actflow model-based analysis complements previous studies characterizing the role of DLPFC in working memory (Cai et al., 2021; Finn et al., 2019; Mencarelli et al., 2019; Senkowski et al., 2022) by revealing influences from default mode and frontoparietal networks that likely drive DLPFC’s involvement in n-back cognitive processes. Further, actflow models allowed us to estimate the degree to which distributed afferent processes—as opposed to recurrent within-region processes—play a role (47% of n-back contrast inter-subject variance explained) in right DLPFC working-memory-related activity. As previously mentioned, this single-step directed actflow model represents a starting point for a future more comprehensive multiple-step neurocognitive explanations of working memory (Christophel et al., 2017). Such an explanation would characterize the chain of effects from stimulus (e.g., n-back task stimuli) to network-based cognitive processes (e.g., neural activity changes from differences in working memory load) to behavior (e.g., motor response).

While the particular methods used here are illustrative of the proposed mechanistic actflow paradigm for explaining neurocognitive activations, there are many opportunities for further improvements. The limitations of the current study can perhaps be best illustrated by contrasting it with an idealized experiment not currently possible due to methodological limitations in neuroscience. Such an ideal study would observe all action potentials and local field potentials throughout the human brain in real time. This would contrast with the present study’s use of fMRI, which has limited spatial resolution (2.0 mm voxels) and temporal resolution (720 ms time points) but whole-brain coverage that matches the ideal. FMRI detects blood-oxygen level dependent changes in MRI signals, which is an indirect measure of aggregate action potentials and local field potentials (Kahn et al., 2013; Logothetis et al., 2001; Shmuel & Leopold, 2008).

The ideal study’s perfect spatial coverage would be essential for reducing potential causal confounds. Given that fMRI has full spatial coverage, this benefit extended to the present study. However, the ideal study’s perfect spatial resolution would allow the FC algorithms used in the present study to even better account for potential causal confounders, such as neural signals lost through averaging into observed voxel time series. The ideal study’s perfect temporal resolution would present a challenge to the FC algorithms used here, since action potentials and downstream changes in local field potentials would occur with a temporal lag. Such lags would provide additional constraints on causal inferences not accounted for in the present study, but which are utilized in actflow studies utilizing higher temporal resolution methods (Mill et al., 2022). As mentioned above, another important component in the ideal study would be to follow the causal chain from stimulus to distributed cognitive processes to behavior. This more comprehensive explanation of the DLPFC n-back effect is beyond the scope of the present study, but something close to this level of explanation—of a different set of neurocognitive phenomena—has been achieved in a recent actflow study (Ito et al., 2022). Finally, the ideal study would use stimulation and lesion approaches to fully verify the causal relevance of observed neural signals.

We estimated FC networks using resting-state data, which has become a standard approach in neuroimaging. However, a recent study demonstrated improvements in actflow predictions when using task-state FC (relative to resting-state FC) (Cole et al., 2021). We nonetheless chose to use resting-state FC here, given improvements in explanatory power with this approach. Specifically, successful actflow predictions using FC from a brain state (rest) other than the state of interest (here the n-back task) provided evidence that our inferences were based on latent connectivity properties that generalize across brain states (McCormick et al., 2022). This approach contrasts with focusing on an FC configuration unique to the current state of interest, with no information regarding the generalizability of the resulting actflow model to other states/tasks. Further, inferring FC networks using data from another state reduces the chance of overfitting to noise (Lever et al., 2016), which could potentially result in false-positive inferences. An additional practical consideration of using resting-state data here, given that our dataset included much more resting-state data than task data, is that FC estimation is substantially improved by including more time points, such that actflow predictions can be improved using the brain state(s) with the greatest amount of data (Cole et al., 2021; Sanchez-Romero & Cole, 2021). Using FC estimates from the state of interest nonetheless provides an opportunity for future research, given the possibility that some details of the actflow mechanisms generating cognitive effects were not observed with the present approach (e.g., task-specific DLPFC FC updates during the n-back task).

In our study, FC networks were inferred using the PC algorithm due to its effective use of statistical conditional independence information and causal principles, implementation simplicity, efficient run times with many variables, and adaptability (here we used linear conditional association tests, but nonlinear or nonparametric tests can also be considered for future studies). Nonetheless, the PC algorithm has two important assumptions, which we were only able to partially address in our version of the algorithm: (i) Acyclicity of the true network. Here we used a PC adjacency discovery step that previously showed good performance on cyclic simulations (Sanchez-Romero et al., 2019), and also showed good control of false-positive connections in our simulations. In addition, we excluded PC orientation rules based specifically on the assumption of acyclicity (see PC algorithm section), preferring to retain undirected connections that may suggest the presence of cycles rather than risking incorrect orientations. Note that we included cycles in our simulations, allowing us to assess how detrimental violations of the non-cyclic assumptions of the PC algorithm are likely to be. (ii) No unmeasured confounders. The presence of unmeasured confounders weakens the mechanistic interpretation of the PC results due to uncertainty about the real causal driver of the inferred connection between region pairs. There are standard strategies to mitigate the impact of confounders in fMRI applications. In our study we followed two: (1) Account for temporal and physiological confounders by detrending and applying nuisance regression to the BOLD time series (see Methods, Ciric et al., 2017; Murphy et al., 2009), and (2) account for all regions of interest by using a brain parcellation with full cortical coverage. In cases when suspected confounders are hard to measure, another strategy is to use directed FC methods that explicitly model the presence of unmeasured confounders like Fast Causal Inference (Malinsky & Spirtes, 2017; Spirtes et al., 2000) and Two-Step (Sanchez-Romero et al., 2019). It is not straightforward to determine how actflow models should be parameterized with these more complex directed FC approaches, but solving this problem is a promising opportunity to strengthen the mechanistic validity of actflow models.

Of particular relevance for empirical applications of actflow is the PC algorithm’s collider orientation step (**Box 1.2.1**), since collider causal structures (*X* → *Z* ← *Y*) are the structures of most interest for inferring the generation of neurocognitive function via actflow. This is because the generation of functionality likely requires multiple elements mixing and interacting in some way—the exact situation described by a collider. This requirement for multiple elements interacting to generate something new is not unique to neural processing, as it is a property of the universe that compositional elements (e.g., hydrogen and oxygen) interact to produce unique configurations with new properties (e.g., H_2_O/water). And so we conceptualize the activity of multiple brain regions interacting as they influence right DLPFC—via a collider causal structure—to help produce working memory functionality. Notably, collider-based interactions may benefit from non-linearities (e.g., super-linear DLPFC activity when two or more inputs are present), suggesting potentially fruitful future research focusing on non-linear collider interactions.

The PC algorithm, as applied here, estimates contemporaneous functional associations from fMRI data, but for neural data acquired with higher temporal resolution, such as electroencephalography (EEG) or magnetoencephalography (MEG), it would be possible to apply temporal FC methods based on autoregressive processes that identify temporally lagged functional interactions between brain regions (Amblard & Michel, 2013; Biswas & Shlizerman, 2022; Gilson et al., 2019; Malinsky & Spirtes, 2018; Moneta et al., 2011; Novelli et al., 2019; Runge, 2018; Shen et al., 2019; Stramaglia et al., 2014). For example, as mentioned above, Mill et al. (2022) recently showed how temporally resolved FC can characterize the dynamics of the task-evoked activity flows that produce cognitive computations. We chose not to use lag-based autoregressive modeling approaches here due to the fundamental sluggishness of the hemodynamic response (∼6 second delay from neural input to peak hemodynamic signal), as well as the low temporal sampling of each neural population with fMRI (720 ms TR here). These temporal properties contrast substantially with the range of direct-interaction lags between neural populations, which have been shown with microstimulation in non-human primates to be on the order of 3-6 ms (Firmin et al., 2014; Yazdan-Shahmorad et al., 2018). While direct-interaction lags may be longer between distant regions within the human brain, they are unlikely to be on the order of 720 ms (or 6 seconds), as would likely be required for effective lag estimation with fMRI data.

The FC methods presented here, despite not considering the temporal information of the BOLD signal, assume stationarity of the signals, in particular related to the possibility that two brain regions *X* and *Y* are statistically associated only because they are confounded by a temporal trend *t* (e.g., *X* ← *t* → *Y*). To mitigate the confounding effect of a potential temporal trend and to better estimate the association between brain regions, we included a linear detrending step in our empirical BOLD signal preprocessing (see Materials and Methods).

In our study we focused on FC methods leveraging conditional independence information and particular causal principles, but other available directed FC methods could also in principle be applied. Promising alternatives are an efficient implementation of the popular dynamical causal modelling (DCM) approach (Frässle et al., 2021), and neural network modeling approaches, such as mesoscale individualized neurodynamic modeling (MINDy) (Singh et al., 2020) and current-based decomposition (CURBD) (Perich et al., 2020). These methods use different causal principles and estimation techniques than the PC algorithm (and related Bayesian network methods (Ramsey et al., 2011, 2017)), and thus offer future opportunities to explore the robustness and diversity of possible mechanistic actflow explanations across diverse FC procedures.

Related studies have presented statistical models to predict task activations using resting-state FC and structural connectivity (Tavor et al., 2016), spatial maps (e.g., default mode network) (Dohmatob et al., 2021), and surface-level time series (Ngo et al., 2022). Importantly, these models differ fundamentally from the actflow approach in that they are optimized for prediction rather than causal mechanism *and* prediction. Thus, while these approaches can reveal the general factors that may contribute to task activations (e.g., structural connectivity patterns), unlike actflow they are not designed to reveal how specifically those factors contribute to the generation of task activations (and associated cognition).

Taken together, the results presented here show that actflow models built using directed FC methods can accurately predict activations for a wide variety of brain regions and task conditions, offering a basis for developing future mechanistic explanations that advance our understanding of network-based distributed cognitive computations in the human brain.

## Acknowledgements

The authors acknowledge support by the US National Institutes of Health under awards R01 AG055556, R01 MH109520 and R01 EB022858-03, and the US National Science Foundation under award 2219323. Data were provided, in part, by the Human Connectome Project, WU-Minn Consortium (Principal Investigators: D. Van Essen and K. Ugurbil; 1U54MH091657) funded by the 16 NIH Institutes and Centers that support the NIH Blueprint for Neuroscience Research; and by the McDonnell Center for Systems Neuroscience at Washington University. The authors acknowledge the Office of Advanced Research Computing (OARC) at Rutgers The State University of New Jersey, for providing access to the Amarel cluster and associated research computing resources that have contributed to the results reported here. The content is solely the responsibility of the authors and does not necessarily represent the official views of any of the funding agencies. The authors also thank Joseph D. Ramsey and Kevin Bui for support with the Tetrad software.

## Notes

### Competing Interest Statement

The authors have declared no competing interest.

### Summary of Updates

We remove incorrect hyperlinks. We add a new Table 1 in Introduction and some new comments in Discussion and Results. Results remained unaltered.

## References

1. Aliferis, C. F., Statnikov, A., Tsamardinos, I., Mani, S., & Koutsoukos, X. D. (2010). Local causal and markov blanket induction for causal discovery and feature selection for classification part i: Algorithms and empirical evaluation. Journal of Machine Learning Research, 11(1). https://jmlr.org/papers/v11/aliferis10a.html

2. Amblard, P.-O., & Michel, O. J. (2013). The relation between Granger causality and directed information theory: A review. Entropy, 15(1), 113–143. https://doi.org/10.3390/e15010113

3. Aquino, K. M., Fulcher, B. D., Parkes, L., Sabaroedin, K., & Fornito, A. (2020). Identifying and removing widespread signal deflections from fMRI data: Rethinking the global signal regression problem. NeuroImage, 212, 116614. https://doi.org/10.1016/j.neuroimage.2020.116614

4. Barch, D. M., Burgess, G. C., Harms, M. P., Petersen, S. E., Schlaggar, B. L., Corbetta, M., Glasser, M. F., Curtiss, S., Dixit, S., & Feldt, C. (2013). Function in the human connectome: Task-fMRI and individual differences in behavior. Neuroimage, 80, 169–189. https://doi.org/10.1016/j.neuroimage.2013.05.033

5. Berkson, J. (1946). Limitations of the application of fourfold table analysis to hospital data. Biometrics Bulletin, 2(3), 47–53. https://www.jstor.org/stable/3002000

6. Bishop, C. M. (2006). Pattern recognition and machine learning. springer. isbn: 978-1-4939-3843-8. https://dl.acm.org/doi/10.5555/1162264

7. Biswas, R., & Shlizerman, E. (2022). Statistical perspective on functional and causal neural connectomics: The Time-Aware PC algorithm. PLOS Computational Biology, 18(11), e1010653. https://doi.org/10.1371/journal.pcbi.1010653

8. Cabeza, R., & Nyberg, L. (2000). Imaging Cognition II: An Empirical Review of 275 PET and fMRI Studies. Journal of Cognitive Neuroscience, 12(1), 1–47. https://doi.org/10.1162/08989290051137585

9. Cai, W., Ryali, S., Pasumarthy, R., Talasila, V., & Menon, V. (2021). Dynamic causal brain circuits during working memory and their functional controllability. Nature Communications, 12(1), 3314. https://doi.org/10.1038/s41467-021-23509-x

10. Carlson, S., Martinkauppi, S., Rämä, P., Salli, E., Korvenoja, A., & Aronen, H. J. (1998). Distribution of cortical activation during visuospatial n-back tasks as revealed by functional magnetic resonance imaging. Cerebral Cortex, 8(8), 743–752. https://doi.org/10.1093/cercor/8.8.743

11. Christophel, T. B., Klink, P. C., Spitzer, B., Roelfsema, P. R., & Haynes, J.-D. (2017). The Distributed Nature of Working Memory. Trends in Cognitive Sciences, 21(2), 111–124. https://doi.org/10.1016/j.tics.2016.12.007

12. Ciric, R., Wolf, D. H., Power, J. D., Roalf, D. R., Baum, G. L., Ruparel, K., Shinohara, R. T., Elliott, M. A., Eickhoff, S. B., Davatzikos, C., Gur, R. C., Gur, R. E., Bassett, D. S., & Satterthwaite, T. D. (2017). Benchmarking of participant-level confound regression strategies for the control of motion artifact in studies of functional connectivity. NeuroImage, 154, 174–187. https://doi.org/10.1016/j.neuroimage.2017.03.020

13. Cole, M. W., Ito, T., Bassett, D. S., & Schultz, D. H. (2016). Activity flow over resting-state networks shapes cognitive task activations. Nature Neuroscience, 19(12), 1718. https://doi.org/10.1038/nn.4406

14. Cole, M. W., Ito, T., Cocuzza, C., & Sanchez-Romero, R. (2021). The functional relevance of task-state functional connectivity. Journal of Neuroscience. https://doi.org/10.1523/JNEUROSCI.1713-20.2021

15. Colenbier, N., Van de Steen, F., Uddin, L. Q., Poldrack, R. A., Calhoun, V. D., & Marinazzo, D. (2020). Disambiguating the role of blood flow and global signal with partial information decomposition. NeuroImage, 213, 116699. https://doi.org/10.1016/j.neuroimage.2020.116699

16. Colombo, D., & Maathuis, M. H. (2014). Order-independent constraint-based causal structure learning. The Journal of Machine Learning Research, 15(1), 3741–3782. https://jmlr.org/papers/v15/colombo14a.html

17. Constantinidis, C., Franowicz, M. N., & Goldman-Rakic, P. S. (2001). The sensory nature of mnemonic representation in the primate prefrontal cortex. Nature Neuroscience, 4(3), 311–316. https://doi.org/10.1038/85179

18. Courtney, S. M., Petit, L., Maisog, J. M., Ungerleider, L. G., & Haxby, J. V. (1998). An Area Specialized for Spatial Working Memory in Human Frontal Cortex. Science, 279(5355), 1347–1351. https://doi.org/10.1126/science.279.5355.1347

19. Curtis, C. E., & D’Esposito, M. (2003). Persistent activity in the prefrontal cortex during working memory. Trends in Cognitive Sciences, 7(9), 415–423. https://doi.org/10.1016/S1364-6613(03)00197-9

20. Dohmatob, E., Richard, H., Pinho, A. L., & Thirion, B. (2021). Brain topography beyond parcellations: Local gradients of functional maps. NeuroImage, 229, 117706. https://doi.org/10.1016/j.neuroimage.2020.117706

21. Eberhardt, F., Glymour, C., & Scheines, R. (2005). On the number of experiments sufficient and in the worst case necessary to identify all causal relations among N variables. Proceedings of the Twenty-First Conference on Uncertainty in Artificial Intelligence, 178–184. https://dl.acm.org/doi/proceedings/10.5555/3020336

22. Finn, E. S., Huber, L., Jangraw, D. C., Molfese, P. J., & Bandettini, P. A. (2019). Layer-dependent activity in human prefrontal cortex during working memory. Nature Neuroscience, 22(10), Article 10. https://doi.org/10.1038/s41593-019-0487-z

23. Firmin, L., Field, P., Maier, M. A., Kraskov, A., Kirkwood, P. A., Nakajima, K., Lemon, R. N., & Glickstein, M. (2014). Axon diameters and conduction velocities in the macaque pyramidal tract. Journal of Neurophysiology, 112(6), 1229– 1240. https://doi.org/10.1152/jn.00720.2013

24. Frässle, S., Harrison, S. J., Heinzle, J., Clementz, B. A., Tamminga, C. A., Sweeney, J. A., Gershon, E. S., Keshavan, M. S., Pearlson, G. D., Powers, A., & Stephan, K. E. (2021). Regression dynamic causal modeling for resting-state fMRI. Human Brain Mapping. https://doi.org/10.1002/hbm.25357

25. Fu, S., & Desmarais, M. C. (2010). Markov blanket based feature selection: A review of past decade. Proceedings of the World Congress on Engineering, 1, 321–328. isbn: 978-988-17012-9-9. https://www.iaeng.org/publication/WCE2010/

26. Gilson, M., Kouvaris, N. E., Deco, G., Mangin, J.-F., Poupon, C., Lefranc, S., Rivière, D., & Zamora-López, G. (2019). Network analysis of whole-brain fMRI dynamics: A new framework based on dynamic communicability. NeuroImage, 201, 116007. https://doi.org/10.1016/j.neuroimage.2019.116007

27. Glasser, M. F., Coalson, T. S., Robinson, E. C., Hacker, C. D., Harwell, J., Yacoub, E., Ugurbil, K., Andersson, J., Beckmann, C. F., & Jenkinson, M. (2016). A multi-modal parcellation of human cerebral cortex. Nature, 536(7615), 171. https://doi.org/10.1038/nature18933

28. Glasser, M. F., Sotiropoulos, S. N., Wilson, J. A., Coalson, T. S., Fischl, B., Andersson, J. L., Xu, J., Jbabdi, S., Webster, M., Polimeni, J. R., Van Essen, D. C., & Jenkinson, M. (2013). The Minimal Preprocessing Pipelines for the Human Connectome Project. NeuroImage, 80, 105–124. https://doi.org/10.1016/j.neuroimage.2013.04.127

29. Gordon, E. M., Breeden, A. L., Bean, S. E., & Vaidya, C. J. (2014). Working memory-related changes in functional connectivity persist beyond task disengagement. Human Brain Mapping, 35(3), 1004–1017. https://doi.org/10.1002/hbm.22230

30. Guyon, I., Aliferis, C., & Elisseeff, A. (2008). Causal feature selection. In H. Liu & H. Motoda (Eds.), Computational methods of feature selection (pp. 63–82). Chapman & Hall/CRC. https://doi.org/10.1201/9781584888796

31. Guyon, I., & Elisseeff, A. (2003). An introduction to variable and feature selection. Journal of Machine Learning Research, 3(Mar), 1157–1182. https://jmlr.org/papers/v3/guyon03a.html

32. Hanson, S. J., & Burr, D. J. (1990). What connectionist models learn: Learning and representation in connectionist networks. Behavioral and Brain Sciences, 13(3), 471–489. https://doi.org/10.1017/S0140525X00079760

33. He, Y., Jia, J., & Yu, B. (2015). Counting and exploring sizes of Markov equivalence classes of directed acyclic graphs. The Journal of Machine Learning Research, 16(1), 2589–2609. https://jmlr.org/papers/v16/he15a.html

34. Hearne, L. J., Mill, R. D., Keane, B. P., Repovš, G., Anticevic, A., & Cole, M. W. (2021). Activity flow underlying abnormalities in brain activations and cognition in schizophrenia. Science Advances, 7(29), eabf2513. https://doi.org/10.1126/sciadv.abf2513

35. Ito, T., Brincat, S. L., Siegel, M., Mill, R. D., He, B. J., Miller, E. K., Rotstein, H. G., & Cole, M. W. (2020). Task-evoked activity quenches neural correlations and variability across cortical areas. PLOS Computational Biology, 16(8), e1007983. https://doi.org/10.1371/journal.pcbi.1007983

36. Ito, T., Hearne, L. J., & Cole, M. W. (2020). A cortical hierarchy of localized and distributed processes revealed via dissociation of task activations, connectivity changes, and intrinsic timescales. NeuroImage, 221, 117141. https://doi.org/10.1016/j.neuroimage.2020.117141

37. Ito, T., Hearne, L., Mill, R., Cocuzza, C., & Cole, M. W. (2020). Discovering the computational relevance of brain network organization. Trends in Cognitive Sciences, 24(1), 25–38. https://doi.org/10.1016/j.tics.2019.10.005

38. Ito, T., Kulkarni, K. R., Schultz, D. H., Mill, R. D., Chen, R. H., Solomyak, L. I., & Cole, M. W. (2017). Cognitive task information is transferred between brain regions via resting-state network topology. Nature Communications, 8(1), 1–14. https://doi.org/10.1038/s41467-017-01000-w

39. Ito, T., Yang, G. R., Laurent, P., Schultz, D. H., & Cole, M. W. (2022). Constructing neural network models from brain data reveals representational transformations linked to adaptive behavior. Nature Communications, 13(1), Article 1. https://doi.org/10.1038/s41467-022-28323-7

40. Ji, J. L., Spronk, M., Kulkarni, K., Repovš, G., Anticevic, A., & Cole, M. W. (2019). Mapping the human brain’s cortical-subcortical functional network organization. NeuroImage, 185, 35–57. https://doi.org/10.1016/j.neuroimage.2018.10.006

41. Jolles, D. D., van Buchem, M. A., Crone, E. A., & Rombouts, S. A. (2013). Functional brain connectivity at rest changes after working memory training. Human Brain Mapping, 34(2), 396–406. https://doi.org/10.1002/hbm.21444

42. Kahn, I., Knoblich, U., Desai, M., Bernstein, J., Graybiel, A. M., Boyden, E. S., Buckner, R. L., & Moore, C. I. (2013). Optogenetic drive of neocortical pyramidal neurons generates fMRI signals that are correlated with spiking activity. Brain Research, 1511, 33–45. https://doi.org/10.1016/j.brainres.2013.03.011

43. Keane, B. P., Barch, D. M., Mill, R. D., Silverstein, S. M., Krekelberg, B., & Cole, M. W. (2021). Brain network mechanisms of visual shape completion. NeuroImage, 236, 118069. https://doi.org/10.1016/j.neuroimage.2021.118069

44. Kiiveri, H., Speed, T. P., & Carlin, J. B. (1984). Recursive causal models. Journal of the Australian Mathematical Society, 36(1), 30–52. https://doi.org/10.1017/S1446788700027312

45. Kriegeskorte, N., Mur, M., & Bandettini, P. A. (2008). Representational similarity analysis-connecting the branches of systems neuroscience. Frontiers in Systems Neuroscience, 2, 4. https://doi.org/10.3389/neuro.06.004.2008

46. Lever, J., Krzywinski, M., & Altman, N. (2016). Model selection and overfitting. Nature Methods, 13(9), Article 9. https://doi.org/10.1038/nmeth.3968

47. Li, J., Kong, R., Liegeois, R., Orban, C., Tan, Y., Sun, N., Holmes, A. J., Sabuncu, M. R., Ge, T., & Yeo, B. T. (2019). Global signal regression strengthens association between resting-state functional connectivity and behavior. NeuroImage, 196, 126–141. https://doi.org/10.1016/j.neuroimage.2019.04.016

48. Liu, T. T., Nalci, A., & Falahpour, M. (2017). The global signal in fMRI: Nuisance or Information? NeuroImage, 150, 213–229. https://doi.org/10.1016/j.neuroimage.2017.02.036

49. Logothetis, N. K., Pauls, J., Augath, M., Trinath, T., & Oeltermann, A. (2001). Neurophysiological investigation of the basis of the fMRI signal. Nature, 412(6843), Article 6843. https://doi.org/10.1038/35084005

50. Malinsky, D., & Spirtes, P. (2017). Estimating bounds on causal effects in high-dimensional and possibly confounded systems. International Journal of Approximate Reasoning, 88, 371–384. https://doi.org/10.1016/j.ijar.2017.06.005

51. Malinsky, D., & Spirtes, P. (2018). Causal structure learning from multivariate time series in settings with unmeasured confounding. Proceedings of 2018 ACM SIGKDD Workshop on Causal Discovery, 23–47. http://proceedings.mlr.press/v92/malinsky18a

52. McCormick, E. M., Arnemann, K. L., Ito, T., Hanson, S. J., & Cole, M. W. (2022). Latent functional connectivity underlying multiple brain states. Network Neuroscience, 6(2), 570–590. https://doi.org/10.1162/netn_a_00234

53. Meek, C. (1995). Causal Inference And Causal Explanation With Background Knowledge. In Proceedings of the Eleventh Conference on Uncertainty in Artificial Intelligence pp. 403–418. https://dl.acm.org/doi/abs/10.5555/2074158.2074204

54. Mencarelli, L., Neri, F., Momi, D., Menardi, A., Rossi, S., Rossi, A., & Santarnecchi, E. (2019). Stimuli, presentation modality, and load-specific brain activity patterns during n-back task. Human Brain Mapping, 40(13), 3810–3831. https://doi.org/10.1002/hbm.24633

55. Mill, R. D., Gordon, B. A., Balota, D. A., & Cole, M. W. (2020). Predicting dysfunctional age-related task activations from resting-state network alterations. Neuroimage, 221, 117167. https://doi.org/10.1016/j.neuroimage.2020.117167

56. Mill, R. D., Hamilton, J. L., Winfield, E. C., Lalta, N., Chen, R. H., & Cole, M. W. (2022). Network modeling of dynamic brain interactions predicts emergence of neural information that supports human cognitive behavior. PLOS Biology, 20(8), e3001686. https://doi.org/10.1371/journal.pbio.3001686

57. Moneta, A., Chlaß, N., Entner, D., & Hoyer, P. (2011). Causal search in structural vector autoregressive models. NIPS Mini-Symposium on Causality in Time Series, 95–114. http://proceedings.mlr.press/v12/moneta11.html

58. Murphy, K., Birn, R. M., Handwerker, D. A., Jones, T. B., & Bandettini, P. A. (2009). The impact of global signal regression on resting state correlations: Are anti-correlated networks introduced? NeuroImage, 44(3), 893–905. https://doi.org/10.1016/j.neuroimage.2008.09.036

59. Murphy, K., & Fox, M. D. (2017). Towards a consensus regarding global signal regression for resting state functional connectivity MRI. Neuroimage, 154, 169–173. https://doi.org/10.1016/j.neuroimage.2016.11.052

60. Ngo, G. H., Khosla, M., Jamison, K., Kuceyeski, A., & Sabuncu, M. R. (2022). Predicting individual task contrasts from resting-state functional connectivity using a surface-based convolutional network. NeuroImage, 248, 118849. https://doi.org/10.1016/j.neuroimage.2021.118849

61. Nichols, T. E., & Holmes, A. P. (2002). Nonparametric permutation tests for functional neuroimaging: A primer with examples. Human Brain Mapping, 15(1), 1–25. https://doi.org/10.1002/hbm.1058

62. Norman, K. A., Polyn, S. M., Detre, G. J., & Haxby, J. V. (2006). Beyond mind-reading: Multi-voxel pattern analysis of fMRI data. Trends in Cognitive Sciences, 10(9), 424–430. https://doi.org/10.1016/j.tics.2006.07.005

63. Novelli, L., Wollstadt, P., Mediano, P., Wibral, M., & Lizier, J. T. (2019). Large-scale directed network inference with multivariate transfer entropy and hierarchical statistical testing. Network Neuroscience, 3(3), 827–847. https://doi.org/10.1162/netn_a_00092

64. Pearl, J. (2000). Causality: Models, reasoning and inference. Cambridge University Press. isbn: 9780521895606

65. Perich, M. G., Arlt, C., Soares, S., Young, M. E., Mosher, C. P., Minxha, J., Carter, E., Rutishauser, U., Rudebeck, P. H., Harvey, C. D., & Rajan, K. (2020). Mi. BioRxiv, 2020.12.18.423348. https://doi.org/10.1101/2020.12.18.423348

66. Petrides, M. (2000). The role of the mid-dorsolateral prefrontal cortex in working memory. Experimental Brain Research, 133(1), 44–54. https://doi.org/10.1007/s002210000399

67. Ramsey, J. D. (2014). A Scalable Conditional Independence Test for Nonlinear, Non-Gaussian Data. *ArXiv:1401.*5031. http://arxiv.org/abs/1401.5031

68. Ramsey, J. D. (2016). Improving accuracy and scalability of the pc algorithm by maximizing p-value. *ArXiv Preprint ArXiv:1610.00378*. https://arxiv.org/abs/1610.00378

69. Ramsey, J. D., Glymour, M., Sanchez-Romero, R., & Glymour, C. (2017). A million variables and more: The Fast Greedy Equivalence Search algorithm for learning high-dimensional graphical causal models, with an application to functional magnetic resonance images. International Journal of Data Science and Analytics, 3(2), 121–129. https://doi.org/10.1007/s41060-016-0032-z

70. Ramsey, J. D., Hanson, S. J., & Glymour, C. (2011). Multi-subject search correctly identifies causal connections and most causal directions in the DCM models of the Smith et al. Simulation study. NeuroImage, 58(3), 838–848. https://doi.org/10.1016/j.neuroimage.2011.06.068

71. Reichenbach, H. (1956). The direction of time (Vol. 65). Univ of California Press. isbn: 978-0486409269

72. Reid, A. T., Headley, D. B., Mill, R. D., Sanchez-Romero, R., Uddin, L. Q., Marinazzo, D., Lurie, D. J., Valdés-Sosa, P. A., Hanson, S. J., Biswal, B. B., Calhoun, V., Poldrack, R. A., & Cole, M. W. (2019). Advancing functional connectivity research from association to causation. Nature Neuroscience, 1–10. https://doi.org/10.1038/s41593-019-0510-4

73. Reinhart, R. M. (2017). Disruption and rescue of interareal theta phase coupling and adaptive behavior. Proceedings of the National Academy of Sciences, 114(43), 11542–11547. https://doi.org/10.1073/pnas.1710257114

74. Rowe, J. B., & Passingham, R. E. (2001). Working Memory for Location and Time: Activity in Prefrontal Area 46 Relates to Selection Rather than Maintenance in Memory. NeuroImage, 14(1), 77–86. https://doi.org/10.1006/nimg.2001.0784

75. Rowe, J. B., Toni, I., Josephs, O., Frackowiak, R. S. J., & Passingham, R. E. (2000). The Prefrontal Cortex: Response Selection or Maintenance Within Working Memory? Science, 288(5471), 1656–1660. https://doi.org/10.1126/science.288.5471.1656

76. Rumelhart, D. E., Hinton, G. E., & McClelland, J. L. (1986). A general framework for parallel distributed processing. Parallel Distributed Processing: Explorations in the Microstructure of Cognition, 1(45–76), 26. https://ieeexplore.ieee.org/document/6302935

77. Runge, J. (2018). Causal network reconstruction from time series: From theoretical assumptions to practical estimation. Chaos: An Interdisciplinary Journal of Nonlinear Science, 28(7), 075310. https://doi.org/10.1063/1.5025050

78. Sanchez-Romero, R., & Cole, M. W. (2021). Combining multiple functional connectivity methods to improve causal inferences. Journal of Cognitive Neuroscience, 33(2), 180–194. https://doi.org/10.1162/jocn_a_01580

79. Sanchez-Romero, R., Ramsey, J. D., Zhang, K., Glymour, M. R., Huang, B., & Glymour, C. (2019). Estimating feedforward and feedback effective connections from fMRI time series: Assessments of statistical methods. Network Neuroscience, 3(2), 274–306. https://doi.org/10.1162/netn_a_00061

80. Saxe, R., Brett, M., & Kanwisher, N. (2006). Divide and conquer: A defense of functional localizers. NeuroImage, 30(4), 1088–1096. https://doi.org/10.1016/j.neuroimage.2005.12.062

81. Senkowski, D., Sobirey, R., Haslacher, D., & Soekadar, S. R. (2022). Boosting working memory: Uncovering the differential effects of tDCS and tACS. Cerebral Cortex Communications, 3(2), tgac018. https://doi.org/10.1093/texcom/tgac018

82. Shen, Y., Giannakis, G. B., & Baingana, B. (2019). Nonlinear structural vector autoregressive models with application to directed brain networks. IEEE Transactions on Signal Processing, 67(20), 5325–5339. https://doi.org/10.1109/TSP.2019.2940122

83. Shmuel, A., & Leopold, D. A. (2008). Neuronal correlates of spontaneous fluctuations in fMRI signals in monkey visual cortex: Implications for functional connectivity at rest. Human Brain Mapping, 29(7), 751–761. https://doi.org/10.1002/hbm.20580

84. Singh, M. F., Braver, T. S., Cole, M. W., & Ching, S. (2020). Estimation and validation of individualized dynamic brain models with resting state fMRI. NeuroImage, 221, 117046. https://doi.org/10.1016/j.neuroimage.2020.117046

85. Smith, S. M., Beckmann, C. F., Andersson, J., Auerbach, E. J., Bijsterbosch, J., Douaud, G., Duff, E., Feinberg, D. A., Griffanti, L., & Harms, M. P. (2013). Resting-state fMRI in the human connectome project. Neuroimage, 80, 144–168. https://doi.org/10.1016/j.neuroimage.2013.05.039

86. Spirtes, P., & Glymour, C. (1991). An algorithm for fast recovery of sparse causal graphs. Social Science Computer Review, 9(1), 62–72. https://doi.org/10.1177/089443939100900106

87. Spirtes, P., Glymour, C. N., & Scheines, R. (2000). Causation, prediction, and search (Second edition). The MIT Press. https://doi.org/10.7551/mitpress/1754.001.0001

88. Sreenivasan, K. K., Curtis, C. E., & D’Esposito, M. (2014). Revisiting the role of persistent neural activity during working memory. Trends in Cognitive Sciences, 18(2), 82–89. https://doi.org/10.1016/j.tics.2013.12.001

89. Stramaglia, S., Cortes, J. M., & Marinazzo, D. (2014). Synergy and redundancy in the Granger causal analysis of dynamical networks. New Journal of Physics, 16(10), 105003. https://iopscience.iop.org/article/10.1088/1367-2630/16/10/105003

90. Tavor, I., Jones, O. P., Mars, R. B., Smith, S. M., Behrens, T. E., & Jbabdi, S. (2016). Task-free MRI predicts individual differences in brain activity during task performance. Science, 352(6282), 216–220. https://doi.org/10.1126/science.aad8127

91. Thomas, M. S. C., & McClelland, J. L. (2008). Connectionist models of cognition. In The Cambridge handbook of computational psychology (pp. 23–58). Cambridge University Press. https://doi.org/10.1017/CBO9780511816772.005

92. Ugurbil, K., Xu, J., Auerbach, E. J., Moeller, S., Vu, A. T., Duarte-Carvajalino, J. M., Lenglet, C., Wu, X., Schmitter, S., & Van de Moortele, P. F. (2013). Pushing spatial and temporal resolution for functional and diffusion MRI in the Human Connectome Project. Neuroimage, 80, 80–104. https://doi.org/10.1016/j.neuroimage.2013.05.012

93. Van Essen, D. C., Smith, S. M., Barch, D. M., Behrens, T. E., Yacoub, E., Ugurbil, K., & Consortium, W.-M. H. (2013). The WU-Minn human connectome project: An overview. Neuroimage, 80, 62–79. https://doi.org/10.1016/j.neuroimage.2013.05.041

94. Vatansever, D., Manktelow, A. E., Sahakian, B. J., Menon, D. K., & Stamatakis, E. A. (2017). Angular default mode network connectivity across working memory load. Human Brain Mapping, 38(1), 41–52. https://doi.org/10.1002/hbm.23341

95. Verma, T., & Pearl, J. (1992). An algorithm for deciding if a set of observed independencies has a causal explanation. Uncertainty in Artificial Intelligence, 323–330. https://doi.org/10.1016/B978-1-4832-8287-9.50049-9

96. Wager, T. D., & Smith, E. E. (2003). Neuroimaging studies of working memory: *Cognitive, Affective*, & Behavioral Neuroscience, 3(4), 255–274. https://doi.org/10.3758/CABN.3.4.255

97. Wang, M., Yang, Y., Wang, C.-J., Gamo, N. J., Jin, L. E., Mazer, J. A., Morrison, J. H., Wang, X.-J., & Arnsten, A. F. T. (2013). NMDA Receptors Subserve Persistent Neuronal Firing during Working Memory in Dorsolateral Prefrontal Cortex. Neuron, 77(4), 736–749. https://doi.org/10.1016/j.neuron.2012.12.032

98. Weichwald, S., & Peters, J. (2021). Causality in Cognitive Neuroscience: Concepts, Challenges, and Distributional Robustness. Journal of Cognitive Neuroscience, 33(2), 226–247. https://doi.org/10.1162/jocn_a_01623

99. Woodward, J. (2005). *Making things happen: A theory of causal explanation*. Oxford university press. https://doi.org/10.1093/0195155270.001.0001

100. Yazdan-Shahmorad, A., Silversmith, D. B., Kharazia, V., & Sabes, P. N. (2018). Targeted cortical reorganization using optogenetics in non-human primates. ELife, 7, e31034. https://doi.org/10.7554/eLife.31034

101. Zhang, K., Peters, J., Janzing, D., & Schölkopf, B. (2011). Kernel-based Conditional Independence Test and Application in Causal Discovery. Proceedings of the Twenty-Seventh Conference on Uncertainty in Artificial Intelligence, 804–813. https://dl.acm.org/doi/10.5555/3020548.3020641

